# Phycobilins as potent food bioactive broad-spectrum inhibitor compounds against M^pro^ and PL^pro^ of SARS-CoV-2 and other coronaviruses: A preliminary Study

**DOI:** 10.1101/2020.11.21.392605

**Authors:** Brahmaiah Pendyala, Ankit Patras, Chandravanu Dash

**Affiliations:** Food Biosciences and Technology Program, Department of Agricultural and Environmental Sciences, Tennessee State University, Nashville, 37209, TN, USA; Meharry Medical College, Nashville, 37208 TN USA

**Author notes:** Co-Corresponding authors: Brahmaiah Pendyala, Ph.D., Research Scientist, Department of Agricultural and Environmental Sciences, Tennessee State University, Nashville, TN 37209, USA, Ankit Patras, Ph.D., Associate Professor, Department of Agricultural and Environmental Sciences, Tennessee State University, Nashville, TN 37209, USA.

**Keywords:** Food bioactive constituents, broad-spectrum inhibitors, coronaviruses, SARS-CoV-2, COVID-19, main protease, papain like protease

## Abstract

In the twenty first century, we have witnessed three corona virus outbreaks; SARS in 2003, MERS in 2012 and ongoing pandemic COVID-19. To prevent outbreaks by novel mutant strains, we need broad-spectrum antiviral agents that are effective against wide array of coronaviruses. In this study, we scientifically investigated potent food bioactive broad-spectrum antiviral compounds by targeting M^pro^ and PL^pro^ proteases of CoVs using *in silico* and *in vitro* approaches. The results revealed that phycocyanobilin (PCB) showed potential inhibitor activity against both proteases. PCB had best binding affinity to M^pro^ and PL^pro^ with IC_50_ values of 71 μm and 62 μm, respectively. In addition, *in silico* studies of M^pro^ and PL^pro^ enzymes of other human and animal CoVs indicated broad spectrum inhibitor activity of the PCB. Like PCB, other phycobilins such as phycourobilin (PUB), Phycoerythrobilin (PEB) and Phycoviolobilin (PVB) showed similar binding affinity to SARS-CoV-2 M^pro^ and PL^pro^

## 1. Introduction

Coronaviruses (CoVs) belongs to the subfamily of *Orthocoronavirinae,* family *Coronavidae,* order *Nidovirales.* They are large (average diameter of 120 nm), enveloped, positive-sense single-stranded RNA viruses with genome size of ~26 to 32 kb (Woo et al., 2010). Based on antigen cross reactivity and genetic makeup, there are four sub-groups (alpha, beta, gamma and delta) which subdivided into 26 different species of coronaviruses (Cleri, Ricketti, & Vernaleo, 2010). CoVs cause diseases in mammals and birds, alpha and beta group CoVs are pathogenic to humans (Paules, Marston, & Fauci, 2020). The seven CoVs that can cause infectious diseases in humans are HCoV-229E, HCoV-NL63, HCoV-OC43, HCoV-HKU1, severe acute respiratory syndrome coronavirus (SARS-CoV), Middle East respiratory virus coronavirus (MERS-CoV) and 2019-nCoV (2019-novel coronavirus) or SARS-CoV-2 (Hamre & Procknow, 1966; McIntosh, Becker, & Chanock, 1967; Van Der Hock et al., 2004; Woo et al., 2005; Drosten et al., 2003; Bermingham et al., 2012; Wu et al., 2020). The first four common CoVs persistently circulate in humans and responsible for 10 to 30 % of common colds (Paules, Marston, & Fauci, 2020). Other three are deadly viruses are etiological agents of fatal respiratory syndromes SARS, MERS and coronavirus disease 2019 (COVID-19) respectively. The SARS epidemic in 2003 ended with 8098 reported cases, 774 deaths (fatality rate 9.7 %), whereas MERS outbreak in 2012 caused 2494 reported cases, 858 deaths (fatality rate 34%) (WHO, 2003; Alfaraj et al., 2019). The COVID-19, current pandemic outbreak first identified in2019, reported > 37.1 million confirmed cases with > 1.07 million deaths (fatality rate 2.9 %) as of 12 October 2020 (WHO, 2020). Avian infectious bronchitis virus (IBV), feline infectious peritonitis virus (FIPV), canine CoV, and porcine transmissible gastroenteritis virus (TGEV) cause respiratory and enteric diseases in farm and domestic pet animals (Cavanagh, 2007; Pederson, 2009; Pratelli, 2006; Odend’hal, 1983).

Till now, there are no approved vaccines and therapeutic drugs for COVID-19 or other human coronavirus infections, and lack of enough clinical trial data to make treatment decisions. Although vaccines have been developed against animal viruses IBV, canine CoV, and TGEV to help prevent serious diseases (Liu & Kong, 2004; Carmichael, 1999; Park et al., 1998), some potential problems such as; recombination events between field and vaccine strains, emergence of novel serotypes, and antibody-dependent enhancement remain. Currently, rapid development of vaccines and repurposing of approved antivirals drugs (for e.g. remdesivir) are major clinical approaches of pandemic preparedness plan. Development of broad-spectrum antiviral agents which are effective against wide range of CoVs and other classes of viruses including emerging ones could be a promising strategy (Fauci, 2006; Bekerman & Einav, 2015; Cho & Glenn, 2020).

Broad-spectrum antiviral targeting can be classified strategies into two categories; i) entry inhibitors interact with existing virus particles outside of cells and prevent infection (Hangartner, Zinkernagel, & Hengartner, 2006) and ii) replication inhibitors aimed at stopping viral genome replication to curtail production of new virus particles (De Clerq, 2004). The S glycoprotein of coronaviruses, the main determinant of host cell attachment and viral entry, is not well conserved between HCoVs (Totura & Bavari, 2019). On the other hand, CoV nonstructural proteins (nsps) are highly conserved components of the coronavirus lifecycle that mediate viral replication (Totura & Bavari, 2019). Literature studies reported the following SARS-CoV-2 nsp targets; Main Protease (M^pro^), Papain-like protease (PL^pro^), Nsp3, RdRp, Helicase, Nsp14, Nsp15, Nsp16, N protein to inhibit virus replication (Wu et al., 2020). Proteolytic processing of viral polyproteins into functional nsps by two viral proteases, the M^pro^ and PL^pro^ is an important event of CoV lifecycle. The M^pro^ acts on minimum 11 cleavage sites of replicase 1ab, ~790 kDa; at recognition sequence Leu-Gln↓(Ser, Ala, Gly) (↓ indicates cleavage site) for most cleavage sites, block viral replication (Zhang et al., 2020). PL^Pro^ enzyme hydrolyses the peptide bond at carboxyl side of glycine (P1 position) and releases nsp1, nsp2, and nsp3 functional proteins, which play key role in viral replication (Rut et al., 2020). Therefore, these proteases would be potential targets for development of broad-spectrum antiviral drugs. The crystal structures of CoVs M^pro^ and PL^pro^ enzymes are available for public access in protein data bank (PDB) (Table 2).

Natural phytochemicals based drugs are gaining importance in the modern world healthcare sector, because of its less toxicity, effective health benefits and its potential use in conjunction with preexisting therapies. Several literature studies have been reported antiviral properties of phytochemicals against CoVs and other viruses (Mani et al., 2020; Ghildiyal et al., 2020). In view of the issues posed above, the identification of natural broad-spectrum antiviral agents against the CoVs is a more reasonable and attractive prospect, could provide an effective first line of defense against future emerging CoVs related diseases. Herein, we report the natural phycobilins as potent natural broad-spectrum inhibitor compounds against 3CLpro and PLpro of SARS-CoV-2 and other CoVs via *in silico* and *in-vitro* approaches.

## 2. Materials and Methods

### 2.1. In silico screening of inhibitor compounds

#### 2.1.1. Preparation of protein and ligand for docking

The crystal structures of M^pro^ (PDB ID – 6LU7) and PL^pro^ of SARS-CoV-2 (PDB ID – 6WUU) and other CoVs (PDB ID numbers were given in Table 2) used in this study were obtained from RCSB protein data bank. Ligand structures were obtained from Pubchem and Chemical Entities of Biological Interest (ChEBI) as SDF format, Open Babel was used for format transformation or 3D coordinate generation for the uploaded files (O’Boyle et al. 2011). The MGLTools were used to delete other chains, heteroatoms (included water), adding missing atoms, hydrogens, charges. The files (pdbqt) were further prepared for proteins and ligands binding.

#### 2.1.2. Molecular docking studies

Autodock Vina was used as docking engine for docking. It is critical to define the docking grid box appropriately due to the characteristic of small molecule docking procedure (Trott and Olson, 2010). The docking box is defined as the center of native ligand coordinates with 40Å×40Å×40Å in length to include the residues of entire cavity, and the exhaustiveness level was set on 12 with number of modes 10. For visualization, the docking results PDBQT files were exported and docked protein-ligand complex structures were visualized using Pymol. Active site residues within 3 or 3.5 Å of ligand and polar contacts were determined with this same tool.

### 2.2. In-vitro enzymatic assays

For enzyme inhibition studies, selected phytochemicals; PCB, Quercetin, Riboflavin, Cyanadin, Daidzein and Genestein were purchased from Santa Cruz Biotechnology (Santa Cruz, CA). Enzyme assay kits; 3CL Protease, MBP-tagged (SARS-CoV-2) assay (Catalog #79955) and papain-like protease (SARS-CoV-2) assay kit: protease activity (Catalog #79995) were purchased from BPS Bioscience (San Diego, CA).

#### 2.2.1. M^pro^ assay

Fluorescence resonance energy transfer (FRET)-based cleavage assay (Zhu et al., 2011) was used for *in-vitro* enzyme inhibition study. SARS-CoV-2 M^pro^ or 3CL Protease, GenBank Accession No. YP_009725301, amino acids 1-306 (full length), with an N-terminal MBP-tag, expressed in an *E. coli* and its fluorescent substrate with cleavage site (indicated by ↓) DABCYL-KTSAVLQ↓SGFRKME-EDANS, inhibitor control (GC376) and the assay buffer composed of 20 mM Tris, 100 mM NaCl, 1 mM EDTA, 1 mM DTT, pH 7.3 were used. Initially, 15 μL of the SARS-CoV-2 M^pro^ in reaction buffer at the final concentration of 10 ng/ μL and 5 μL of inhibitor control (GC376, final conc. 50 μM)/ test inhibitor (10 – 600 μM) / inhibitor solvent (positive control) was pipetted into a 384-well plate. Stock solutions of the compounds were prepared with 100% DMSO. Afterwards, the plate was pre-incubated for 30 min at room temperature with slow shaking. Then the enzymatic reaction was initiated by addition of 5 μL of the substrate dissolved in the reaction buffer to 25 μL final volume (final conc. 50 μM), incubated at room temperature for 4 h. The fluorescence signal of the Edans generated due to the cleavage of the substrate by the Mpro was monitored at an excitation at 360 nm with emission wavelength of 460 nm, using a spectrophotometric microplate reader (Synergy H1 Hybrid Multi-Mode Reader; BioTek Instruments, Inc., Winooski, VT).

#### 2.2.2. PLpro assay

SARS-CoV-2 PL^pro^ papain-like protease, GenBank Accession No. QHD43415, aminoacids 1564-1882, with N-terminal His-tag, expressed in an *E. coli* and its fluorescent substrate Z-Arg-Leu-Arg-Gly-Gly-AMC, inhibitor control (GRL0617) and the assay buffer (40 mM Tris pH 8, 110 mM NaCl, 1 mM DTT) was used for the inhibition assay. Briefly, 30 μL of the SARS-CoV-2 PL^pro^ in reaction buffer at the final concentration of 0.44 ng/ μL and 10 μL of inhibitor control (GRL0617, final conc. 100 μM)/ test inhibitor (10 – 600 μM) / inhibitor solvent (positive control) was pipetted into a 96-well plate. Afterwards, the plate was pre-incubated for 30 min at room temperature with slow shaking. Then the enzymatic reaction was initiated by addition of 10 μL of the substrate dissolved in the reaction buffer to 50 μL final volume (final conc. 25 μM), incubated at room temperature for 40-60 min. The fluorescence signal of the substrate after enzymatic reaction was monitored at an excitation at 360 nm with emission wavelength of 460 nm, using a spectrophotometric microplate reader (Synergy H1 Hybrid Multi-Mode Reader; BioTek Instruments, Inc., Winooski, VT). Triplicate experiments (N = 3) were performed for both M^pro^ and PL^pro^ assays, and the mean value was presented with ± standard deviation (SD).

## 3. Results and Discussion

### 3.1. Selection of phytochemicals for the study

A total of fifteen phytochemicals from different chemical classes were selected based on the previous reports of their potent anti-viral effects (Table S1); linear tetrapyrrole – phycocyanobilin (PCB), flavonols – quercetin, catechin, flavin – riboflavin, anthocyanin – cyanidin, isoflavones – daidzein, genistein, stilbenoid phenol – resveratrol, linear diarylheptanoid – curcumin, Xanthophyll – astaxanthin, carotenes – β-carotene, phenolic alkaloid – capsaicin, phenolic ketone – gingerol, phenolic aldehyde – vanillin, allylbenzene – eugenol, monoterpenoid phenol – thymol.

### 3.2. In silico binding interaction studies of selected phytochemical compounds with SARS-CoV-2 M^pro^ and PL^pro^

The fifteen selected phytochemicals were docked into the active site pocket of SARS-CoV-2 M^pro^ and PL^pro^. Table 1 depicts source, docking score and polar contacts of selected phytochemical bioactive compounds with binding site amino acid residues of SARS-CoV-2 proteases. For M^pro^, the results show phycocyanobilin (PCB) docked with best score or binding energy of −8.6 Kcal/mol followed by Riboflavin (−7.9 Kcal/mol), Cyanidin (−7.9 Kcal/mol), Daidgein (−7.8 Kcal/mol) and Genistein (−7.6 Kcal/mol). Twelve key active site amino acid residues (Tyr 54, Gly 143, His 163, Asp 187, Gln 189, Glu 166, Cys 145, Leu 141, Ser 144, Thr 26, Gln 192, Thr 190) of SARS-CoV-2 M^pro^ involved in polar interactions at distance of ≤ 3 Å with ligand phytochemical compounds. Specific polar contacts of each phytochemical compound were shown in Table 1. In case of PL^pro^, as the reported peptide inhibitor VIR250 is bound to the interface of dimer in the crystal structure of 6WUU (Rut et al., 2020), the docking studies were performed with dimer form. Similarly, PCB docked with best score or binding energy of −9.8 Kcal/mol followed by Astaxanthin, (−9.3 Kcal/mol), β-carotene (−9.2 Kcal/mol), Daidzein (−8.9 Kcal/mol), Riboflavin (−8.5 Kcal/mol) and Genistein (−8.3 Kcal/mol). Eleven key active site aminoacid residues (Asp 164, Tyr 264, Gln 269, Arg 166, Tyr 273, Glu 161, Tyr 268, Lys 157, Leu 162, Gly 266, Ser 170) in chain A and thirteen amino acid residues (Arg 166, Gln 174, Met 208, Glu 161, Glu 167, Cys 155, Lys 232, Met 206, Arg 183, Glu 203, Tyr 268, Tyr 273, Thr 301) in chain B of SARS-CoV-2 PL^pro^ involved in polar interactions at distance of ≤ 3 Å with ligand phytochemical compounds. Specific polar contacts of each phytochemical compound were given in Table 1, and Figure 1 show 3 D representation of binding pocket of M^pro^ and PL^pro^ with top score model pose of PCB.

**Table 1:** Molecular docking results of food bioactive compounds with COVID-19 main protease (M^pro^), papain-like protease (PL^pro^)

| Source | Compounds | M<sup>pro</sup> |  | PL<sup>pro</sup> |  |
| --- | --- | --- | --- | --- | --- |
|  |  | Dock score | Polar contacts | Dock score | Polar contacts |
| Cyanobacteria | Phycocyanobilin | -8.6 | Y54, G143, H163, D187, Q189 | -9.8 | D164 (A), R166 (B), Y264 (A) |
| Fruits, vegetables, seeds & grains | Quercetin | -7.8 | Y54, Q189 | -8 | R166 (B), Q269 (A) |
| Eggs, meat, fruits and vegetables | Riboflavin | -7.9 | E166, C145, H163, L141, S144 | -8.5 | R166 (A), Y264 (A), Y273 (A) |
| Grapes and berries | Cyanidin | -7.8 | S144, H163 | -7.9 | E161 (A), Y268 (A) |
| Legumes | Daidzein | -7.8 | T26, E166, Q192, T190 | -8.9 | K157 (A), D164 (A), R166 (B), Q174 (B) |
| Legumes | Genistein | -7.6 | E166 | -8.3 | K157 (A), L162 (A), Q174 (B), M208 (B) |
| Green tea | Catechin | -7.3 | L141, H163 | -7.1 | E161 (B), R166(A) |
| Grapes and berries skin | Resveratrol | -7 | L141, H163, D187 | -7.2 | R166 (A), E167 (B), C155 (B) |
| Turmeric | Curcumin | -7 | G143, S144, C145 | -8 | K157 (A), K232 (B), Y264 (A) |
| Microalgae | Astaxanthin | -7 | None | -9.3 | G266 (A), M206 (B) |
| Fruits and Vegetables | β-carotene | -6.5 | None | -9.2 | None |
| Chili Pepper | Capsaicin | -6.3 | E166, T190, Q192 | -6.5 | K157 (A), M208 (B) |
| Ginger | Gingerol | -6.1 | G143, S144, C145, H163, E166 | -6.4 | R183 (B), E203 (B), R183 (B) |
| Vanilla | Vanillin | -5 | G143, S144, C145, H163, E166 | -5.4 | Y268 (B), Y273 (B), T301 (B) |
| Cloves | Eugenol | -4.9 | L141, G143, S144, C145, H163 | -5.6 | S170 (A), C155 (B) |
| Thyme | Thymol | -4.8 | None | -5.4 | E203 (B) |

**Table 2:** Molecular docking results of phycocyanobilin with proteases of other pathogenic human and animal CoVs

| CoVs | PDB ID | Dock score | Polar contacts |
| --- | --- | --- | --- |
| **M<sup>pro</sup>** |  |  |  |
| SARS-CoV-1 | 1WOF | -8.5 | Y54, N142, G143, S144, T190 |
| MERS-CoV | 5C3N | -9.3 | H41 (2), Q167, K191 (2), Q195 (2) |
| MHV | 6JIJ | -8.4 | F138, H161, E164, Q187, Q190 |
| TGEV | 2AMP | -8.3 | V26, H41, H162 |
| FIPV | 5EU8 | -8.5 | H41, T47, H162, H163, G167, Q191 |
| IBV | 2Q6F | -8.9 | F46, G141, A142, C143, E187, Q190 |
| HCoV 229 E | 3DBP | -8.3 | I140, H162, E165, G167 |
| HCoV NL63 | 5DWY | -9 | Y53, G142 (2), A143, H163, Q164 |
| HCoV HKU1 | 3D23 | -8.4 | E166 (2), S168 |
| **PL<sup>pro</sup>** |  |  |  |
| SARS-CoV-1 (dimer) | 2FE8 | -8.9 | K158 (A), D165 (A), E168 (A), H172 (B) |
| SARS-CoV-1 (monomer) | 2FE8 | -7.6 | L163, G164, Y269, T302 |
| SARS-CoV-2 (monomer) | 6LU7 | -8.0 | R166, G266 |
| MERS-CoV (monomer) | 4RNA | -8.5 | D164, D165, G248, G277, Y279 |
| TGEV (monomer) | 3MP2 | -8.1 | D80, H153, Q180, G182, Y184 |
| IBV (monomer) | 4X2Z | -7.8 | D150, F151 (2), S152, D153 |

**Figure 1:**
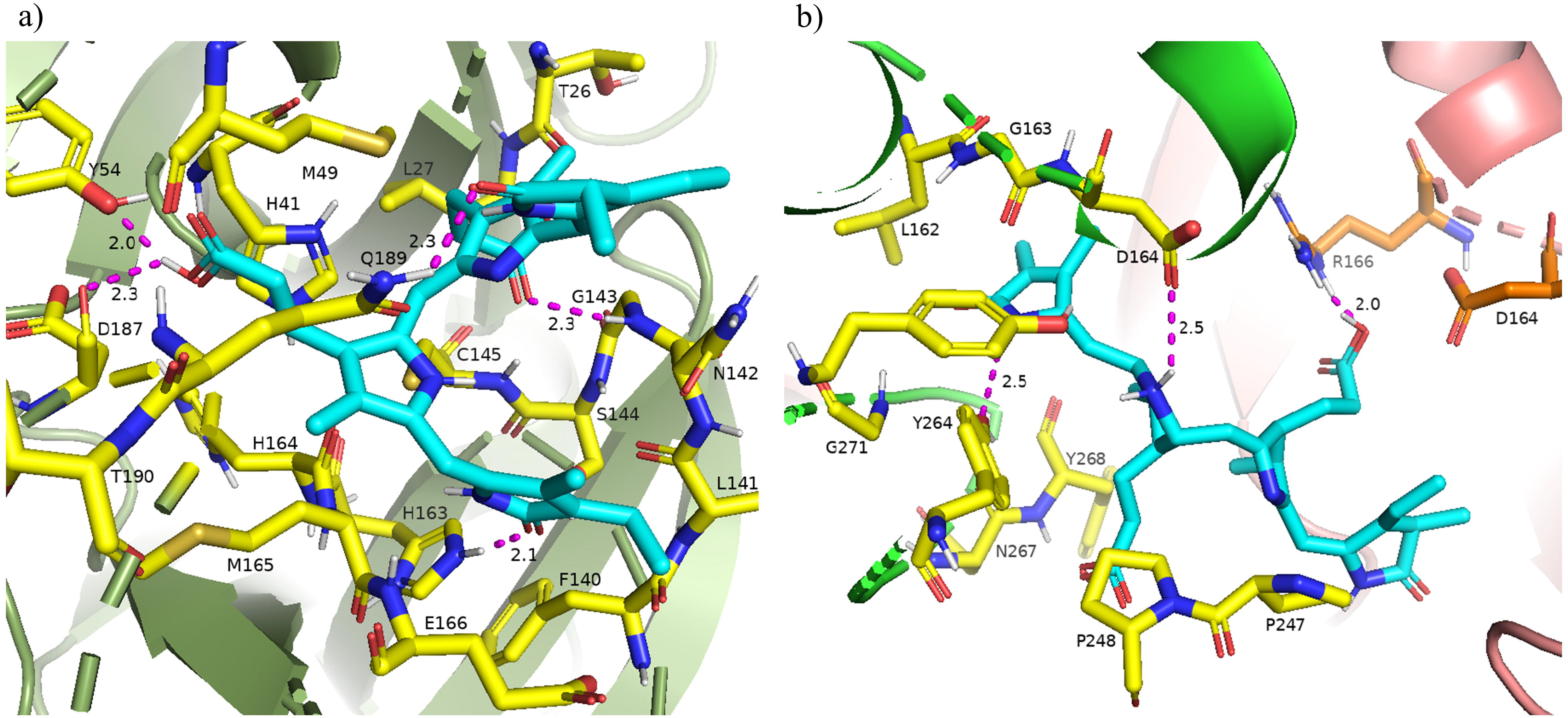
a) 3D binding pocket of SARS-CoV-2 M^pro^ with top model PCB (cyan color), surrounding active site amino acid residues (yellow color) with in 3 Å, remaining residues are represented as a cartoon; b) 3D binding pocket of SARS-CoV-2 PL^pro^ with top model PCB (cyan color), surrounding active site amino acid residues (chain A – yellow color, chain B – orange color) with in 3 Å, remaining residues are represented as a cartoon (chain A – green color, chain B – light pink color). Polar interactions are represented as magenta color.

### 3.3. In vitro enzymatic assay studies to screen potent phytochemical inhibitor compounds against SARS-CoV-2 M^pro^ and PL^pro^

To validate the molecular docking studies, *in-vitro* enzymatic studies were conducted. A positive control without the inhibitor compound in the reaction mixture, an inhibitor control which contains authentic inhibitors GC376 (for M^pro^), GRL0617 (for PL^pro^) were used in this study. The relative activity of the enzymes in the presence of inhibitors was estimated by considering positive control activity as 100 %. Based on *in silico* studies, top six phytochemicals (PCB, quercetin, riboflavin, cyanadin, daidzein and genestein) which showed significant inhibition were selected for M^pro^ enzymatic assay studies. Initial screening results revealed that PCB had higher inhibitor activity followed by quercetin, genestein, cyanidin and riboflavin (p < 0.05) (Fig. S1). Further IC_50_ value of top two compounds; PCB and quercetin was determined, the results show effective IC_50_ value of 71 μM for PCB (Fig. 2) than quercetin (145 μM). For PL^pro^, four compounds (Phycocyanobilin, Riboflavin, Genestein and Quercetin), were screened for inhibitor activity. It was envisaged that PCB showed potent inhibitor activity in comparison to other compounds (Fig. S1), with IC_50_ value of 62 μM (Fig. 2). Over all *in silico* docking and *in vitro* enzyme inhibitor activity results showed PCB as a potent inhibitor against SARS-CoV-2 M^pro^ and PL^pro^. In addition to above mentioned studies (Table S1), other studies reported antiviral activities of herbal and medicinal some natural medicines and plant based bioactives compounds against various virus strains including coronavirus; SARS-CoV-1, herpes simplex, human immunodeficiency virus, hepatitis B and C viruses (Xian et al., 2020). From past few decades, there are numerous efforts to reveal the specific mechanisms of antiviral activities of these natural compounds that effects viral life cycle activities of viral entry, replication, assembly and release (Xian et al., 2020).

**Figure 2:**
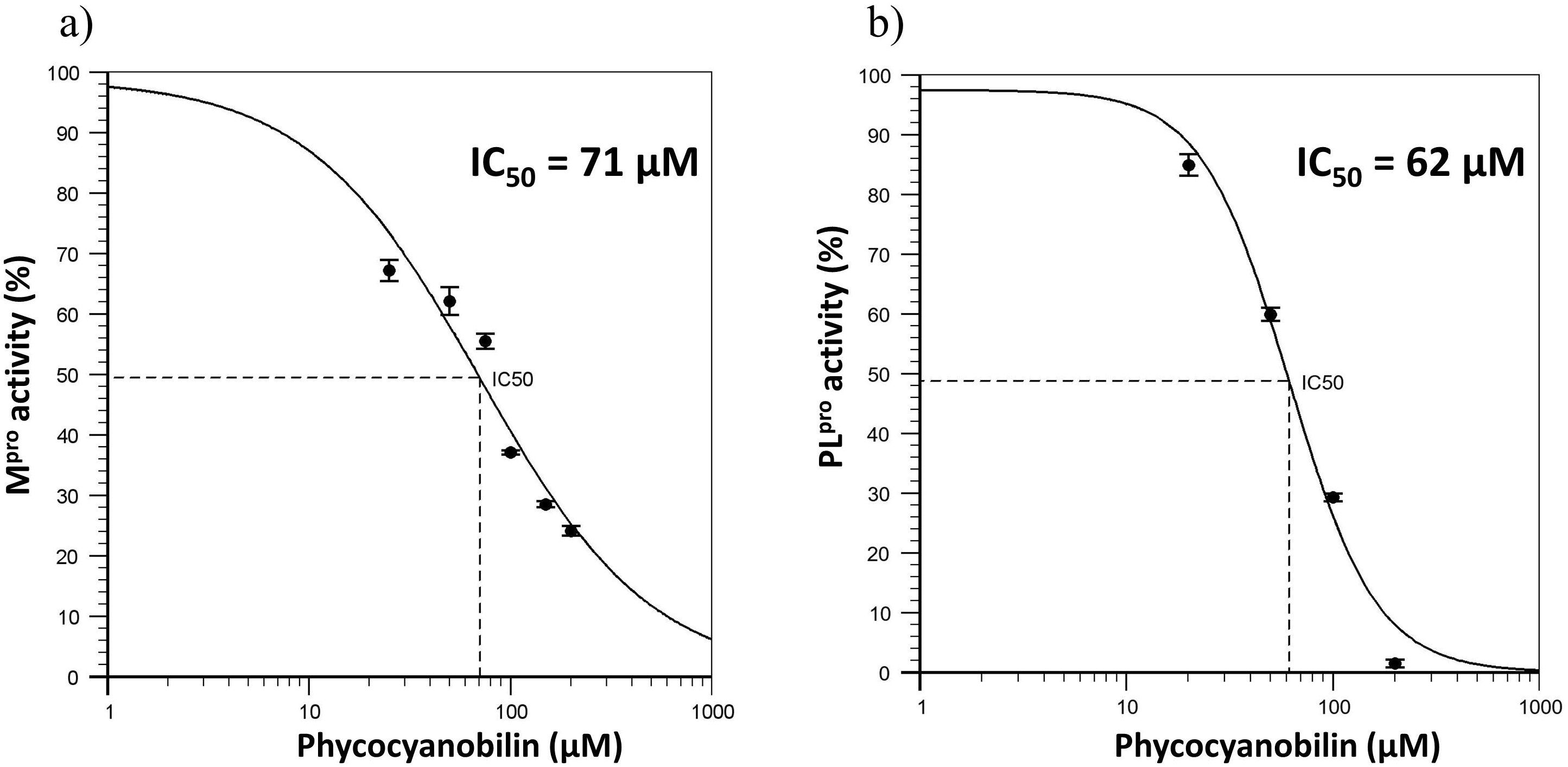
a) Dose response curve of Phycocyanobilin versus M^pro^ activity; b) Dose response curve of Phycocyanobilin versus PL^pro^ activity

### 3.4 Broad-spectrum inhibitor activity of PCB against M^pro^ and PL^pro^

The broad-spectrum efficacy of PCB against CoVs was evaluated by molecular docking studies with available crystal protein data bank (PDB) structure of various human and animal CoVs. Table 2 show the PDB identification code and top docking scores of PCB with M^pro^ and PL^pro^ enzymes of human and animal CoVs. Due to limitation on availability of crystal PDB structures of PL^pro^, both dimer and monomeric forms were used in docking studies. For M^pro^, docking scores are in the range of −8.3 to −9.3 K.cal/mol. PCB showed higher binding affinity with docking score (−9.3 K.cal/mol) for MERS M^pro^ followed by HCoV NL63 (−9.0 K.cal/mol) and IBV (−8.9 K.cal/mol). For PL^pro^, docking scores were in the range of −8.9 to −7.6 K.cal/mol. The results revealed that PCB had higher binding affinity to dimer form of PL^pro^ enzymes than monomeric forms. When compared monomers only, PCB had best docking score for MERS-CoV (−8.5 K.cal/mol) followed by TGEV (−8.1 K.cal/mol) and SARS-CoV-2 (−8.0 K.cal/mol). Figure S2 and S3 show polar contacts of PCB with binding pocket key amino acid residues of M^pro^ and PL^pro^ enzymes of human and animal CoVs. Surprisingly the docking results suggest PCB as a promising broad-spectrum food bioactive inhibitor compound against CoVs proteases. The computed physical properties of phycocyanobilin show rotatable bond count of 10, hydrogen bond donor count of 5 and hydrogen bond acceptor count of 7 (NCBI, 2020), which makes multiple hydrogen bond interactions between the compound and specific amino acid residues located at the active site of the pocket of the protease enzymes. Molecular docking studies indicated that propionic carboxyl and lactam ring carbonyl oxygens of PCB are involved in polar interactions with aminoacid residues of proteases.

### 3.4 Inhibitor activities of other phycobilins

Phycobilins are linear tetrapyrrole chromophore compounds found in certain photosynthetic organisms (cyanobacteria, red algae, glaucophytes, and some cryptomonads) and covalently linked to phycobiliproteins (Beale, 1993). Four types of phycobilins are identified; i) Phycoerythrobilin (PEB), ii) Phycourobilin (PUB), iii) Phycoviolobilin (PVB) and iv) Phycocyanobilin (PCB). Based on results of PCB, the other phycobilins inhibitor activity against SARS-CoV-2 proteases via molecular docking approach was demonstrated and docking scores, polar contacts were given in Table 3. All phycobilins showed strong binding affinity to key amino acids of M^pro^ and PL^pro^ binding pockets. The docking scores were in the order of PUB (−8.7 Kcal/mol) > PCB (−8.6 Kcal/mol) > PEB (−8.2 Kcal/mol) > PVB (−8.2 Kcal/mol) for M^pro^, whereas in case of PL^pro^, the order was PCB (−9.8 Kcal/mol) = PEB (−9.8 Kcal/mol) > PUB (−9.6 Kcal/mol) > PVB (−9.5 Kcal/mol). Nine key binding pocket amino acids (Y54, L141, G143, S144, C145, H163, E166, D187, Q189) of M^pro^ were participated in polar contacts with phycobilins, specific polar contacts of each phycobilin were shown in Figure S4. Ten key binding pocket amino acids (D164 (A), Y264 (A), R166 (A), G266 (A), E161 (A), L162 (A), G271 (A), K232 (A), R166 (B), T301 (B)) of PL^pro^ were participated in polar contacts with phycobilins, specific polar contacts were shown in Figure S5.

**Table 3:**
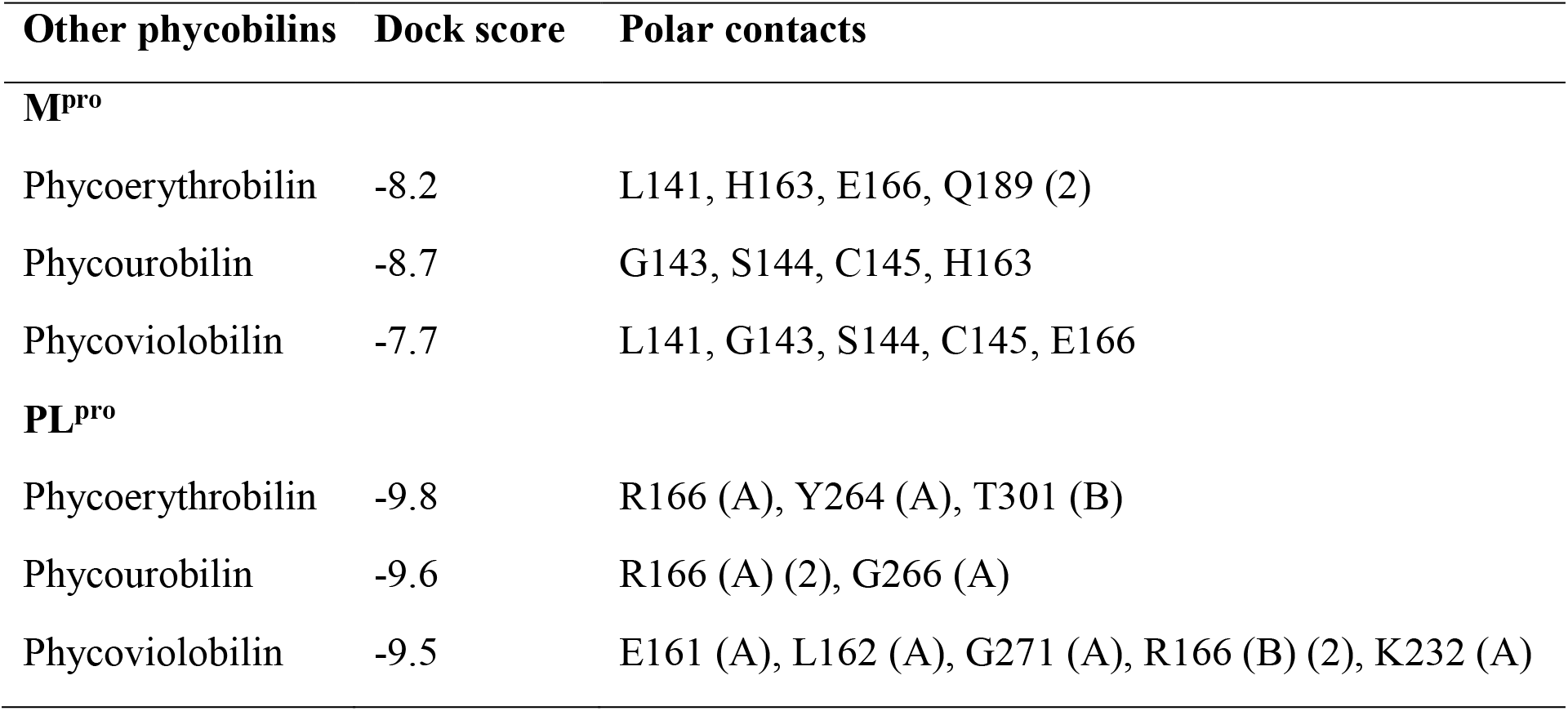
Molecular docking results of other phycobilins with proteases of SARS-CoV-2

| Other phycobilins | Dock score | Polar contacts |
| --- | --- | --- |
| **M<sup>pro</sup>** |  |  |
| Phycoerythrobilin | -8.2 | L141, H163, E166, Q189 (2) |
| Phycourobilin | -8.7 | G143, S144, C145, H163 |
| Phycoviolobilin | -7.7 | L141, G143, S144, C145, E166 |
| **PL<sup>pro</sup>** |  |  |
| Phycoerythrobilin | -9.8 | R166 (A), Y264 (A), T301 (B) |
| Phycourobilin | -9.6 | R166 (A) (2), G266 (A) |
| Phycoviolobilin | -9.5 | E161 (A), L162 (A), G271 (A), R166 (B) (2), K232 (A) |

In addition, potent therapeutic properties such as; peroxy radical scavenging, inhibition of cancer cell proliferation and platelet aggregation, had been reported for phycobilins (Wantanabe, Yabuta & Bito, 2014). Shih et al. (2003) reported direct antiviral activity of phycocyanin against enterovirus 71 in human rhabdomyosarcoma cells and African green monkey kidney cells. Phycobilin compounds can be directly administered orally as phycobiliproteins (complex of phycobilins and protein). For instance, when phycocyanin administered orally to humans, it can be digested and free phycocyanobilin released in the gastrointestinal tract (Wantanabe, Yabuta & Bito, 2014). Thus, noticed therapeutic properties of phycobiliproteins may reflect the effects of their phycobilins (chromophores).

In conclusion, by using, in silico molecular docking, *invitro* enzymatic assay screenings, we discovered PCB as potent phytochemical inhibitors to M^pro^ and PL^pro^ proteases of SARS-CoV-2. Phycocyanobilin had IC_50_ values of 71 and 62 μM for SARS-CoV-2 M^pro^ and PL^pro^, respectively. Further PCB docking studies with other CoVs M^pro^ and PL^pro^ proteases revealed its broad-spectrum inhibitor activity. Also, similar binding affinity of other phycobilins (PEB, PUB and PVB) to these proteases was observed. However *invitro* enzymatic studies with M^pro^ and PL^pro^ of other CoVs consequently *in-vivo* studies on inhibition of CoVs infectivity using human cells and animal models are needed. Further structure guided development of phycobilin lead compounds could rapidly lead to the discovery of a single agent with clinical potential against existing and possible future emerging CoV-associated diseases.

## Note

There are no conflicts to declare.

## References

Alfaraj, S. H., Al-Tawfiq, J. A., Assiri, A. Y., Alzahrani, N. A., Alanazi, A. A., & Memish, Z. A. (2019). Clinical predictors of mortality of Middle East Respiratory Syndrome Coronavirus (MERS-CoV) infection: A cohort study. Travel medicine and infectious disease, 29, 48–50.

Beale, S. I. (1993). Biosynthesis of phycobilins. Chemical Reviews, 93(2), 785–802.

Bekerman, E., & Einav, S. (2015). Combating emerging viral threats. Science, 348(6232), 282–283.

Bermingham, A., Chand, M. A., Brown, C. S., Aarons, E., Tong, C., Langrish, C., … & Pebody, R. G. (2012). Severe respiratory illness caused by a novel coronavirus, in a patient transferred to the United Kingdom from the Middle East, September 2012. Eurosurveillance, 17(40), 20290.

Carmichael, L. E. (1999). Canine viral vaccines at a turning point—a personal perspective. Advances in veterinary medicine, 41, 289.

Cavanagh, D. (2007). Coronavirus avian infectious bronchitis virus. Veterinary research, 38(2), 281–297.

Pedersen, N. C. (2009). A review of feline infectious peritonitis virus infection: 1963-2008. Journal of feline medicine and surgery, 11(4), 225–258.

Cho, N. J., & Glenn, J. S. (2020). Materials science approaches in the development of broad-spectrum antiviral therapies. Nature Materials, 1–4.

Cleri, D. J., Ricketti, A. J., & Vernaleo, J. R. (2010). Severe acute respiratory syndrome (SARS). Infectious Disease Clinics, 24(1), 175–202.

De Clercq, E. (2004). Antivirals and antiviral strategies. Nature Reviews Microbiology, 2(9), 704–720.

Drosten, C., Günther, S., Preiser, W., Van Der Werf, S., Brodt, H. R., Becker, S., … & Berger, A. (2003). Identification of a novel coronavirus in patients with severe acute respiratory syndrome. New England journal of medicine, 348(20), 1967–1976.

Fauci, A. S., & Morens, D. M. (2016). Zika virus in the Americas—yet another arbovirus threat. New England journal of medicine, 374(7), 601–604.

Ghildiyal, R., Prakash, V., Chaudhary, V. K., Gupta, V., & Gabrani, R. (2020). Phytochemicals as Antiviral Agents: Recent Updates. In Plant-derived Bioactives (pp. 279–295). Springer, Singapore.

Hamre, D., & Procknow, J. J. (1966). A new virus isolated from the human respiratory tract. Proceedings of the Society for Experimental Biology and Medicine, 121(1), 190–190.

Hangartner, L., Zinkernagel, R. M., & Hengartner, H. (2006). Antiviral antibody responses: the two extremes of a wide spectrum. Nature Reviews Immunology, 6(3), 231–243.

Liu, S., & Kong, X. (2004). A new genotype of nephropathogenic infectious bronchitis virus circulating in vaccinated and non-vaccinated flocks in China. Avian Pathology, 33(3), 321–327.

Mani, J. S., Johnson, J. B., Steel, J. C., Broszczak, D. A., Neilsen, P. M., Walsh, K. B., & Naiker, M. (2020). Natural product-derived phytochemicals as potential agents against coronaviruses: a review. Virus Research, 197989.

McIntosh, K., Becker, W. B., & Chanock, R. M. (1967). Growth in suckling-mouse brain of” IBV-like” viruses from patients with upper respiratory tract disease. Proceedings of the National Academy of Sciences of the United States of America, 58(6), 2268.

National Center for Biotechnology Information (2020). PubChem Compound Summary for CID 365902, Phycocyanobilin. Retrieved November 9, 2020 from https://pubchem.ncbi.nlm.nih.gov/compound/Phycocyanobilin.

O’Boyle, N. M., Banck, M., James, C. A., Morley, C., Vandermeersch, T., & Hutchison, G. R. (2011). Open Babel: An open chemical toolbox. Journal of cheminformatics, 3(1), 33.

ODEND’HAL, S. T. E. W. A. R. T. (1983). Porcine Transmissible Gastroenteritis Virus. The Geographical Distribution of Animal Viral Diseases, 329.

Park, S., Sestak, K., Hodgins, D. C., Shoup, D. I., Ward, L. A., Jackwood, D. J., & Saif, L. J. (1998). Immune response of sows vaccinated with attenuated transmissible gastroenteritis virus (TGEV) and recombinant TGEV spike protein vaccines and protection of their suckling pigs against virulent TGEV challenge exposure. American journal of veterinary research, 59(8).

Paules, C. I., Marston, H. D., & Fauci, A. S. (2020). Coronavirus infections—more than just the common cold. Jama, 323(8), 707–708.

Pedersen, N. C. (2009). A review of feline infectious peritonitis virus infection: 1963 2008. Journal of feline medicine and surgery, 11(4), 225–225.

Pratelli, A. (2006). Genetic evolution of canine coronavirus and recent advances in prophylaxis. Veterinary Research, 37(2), 191–200.

Rut, W., Lv, Z., Zmudzinski, M., Patchett, S., Nayak, D., Snipas, S. J., … & Olsen, S. K. (2020). Activity profiling and crystal structures of inhibitor-bound SARS-CoV-2 papain-like protease: A framework for anti-COVID-19 drug design. Science Advances, 6(42), eabd4596.

Shih, S. R., Tsai, K. N., Li, Y. S., Chueh, C. C., & Chan, E. C. (2003). Inhibition of enterovirus 71-induced apoptosis by allophycocyanin isolated from a blue-green alga Spirulina platensis. Journal of medical virology, 70(1), 119–125.

Song, J. M., Lee, K. H., & Seong, B. L. (2005). Antiviral effect of catechins in green tea on influenza virus. Antiviral research, 68(2), 66–74.

Totura, A. L., & Bavari, S. (2019). Broad-spectrum coronavirus antiviral drug discovery. Expert opinion on drug discovery, 14(4), 397–397.

Trott, O., & Olson, A. J. (2010). AutoDock Vina: improving the speed and accuracy of docking with a new scoring function, efficient optimization, and multithreading. Journal of computational chemistry, 31(2), 455–461.

Van Der Hoek, L., Pyrc, K., Jebbink, M. F., Vermeulen-Oost, W., Berkhout, R. J., Wolthers, K. C., … & Berkhout, B. (2004). Identification of a new human coronavirus. Nature medicine, 10(4), 368–368.

Watanabe, F., Yabuta, Y., & Bito, T. (2014). Tetrapyrrole compounds of cyanobacteria. In Studies in Natural Products Chemistry (Vol. 42, pp. 341–351). Elsevier.

Woo PC, Huang Y, Lau SK, Yuen KY (August 2010). “Coronavirus genomics andbioinformatics analysis”. Viruses. 2(8): 1804–1804.

Woo, P. C., Lau, S. K., Chu, C. M., Chan, K. H., Tsoi, H. W., Huang, Y., … & Poon, L. L. (2005). Characterization and complete genome sequence of a novel coronavirus, coronavirus HKU1, from patients with pneumonia. Journal of virology, 79(2), 884–895.

World Health Organization. (2003). Consensus document on the epidemiology of severe acute respiratory syndrome (SARS) (No. WHO/CDS/CSR/GAR/2003.11). World Health Organization.

World Health Organization. (2020). Coronavirus disease 2019 (COVID-19): Weekly Epidemiological Update 12 October 2020. Available from: https://www.who.int/docs/default-source/coronaviruse/situation-reports/20201012-weekly-epi-update-9.pdf.

Wu, C., Liu, Y., Yang, Y., Zhang, P., Zhong, W., Wang, Y., … & Zheng, M. (2020). Analysis of therapeutic targets for SARS-CoV-2 and discovery of potential drugs by computational methods. Acta Pharmaceutica Sinica B.

Wu, F., Zhao, S., Yu, B., Chen, Y. M., Wang, W., Song, Z. G., … & Yuan, M. L. (2020). A new coronavirus associated with human respiratory disease in China. Nature, 579(7798), 265–265.

Xian, Y., Zhang, J., Bian, Z., Zhou, H., Zhang, Z., Lin, Z., & Xu, H. (2020). Bioactive natural compounds against human coronaviruses: a review and perspective. Acta Pharmaceutica Sinica B.

Zhang, L., Lin, D., Sun, X., Curth, U., Drosten, C., Sauerhering, L., … & Hilgenfeld, R. (2020). Crystal structure of SARS-CoV-2 main protease provides a basis for design of improved α-ketoamide inhibitors. Science, 368(6489), 409–412.

Zhu, L., George, S., Schmidt, M. F., Al-Gharabli, S. I., Rademann, J., & Hilgenfeld, R. (2011). Peptide aldehyde inhibitors challenge the substrate specificity of the SARS-coronavirus main protease. Antiviral research, 92(2), 204–212.

