## Supplementary information for "Phycobilins as potent food bioactive broad-spectrum inhibitor compounds against M^pro^ and PL^pro^ of SARS-CoV-2 and other coronaviruses: A preliminary Study"

**Table S1:** Antiviral properties of selected food bioactive constituents

| **Bioactive compound** | **Target Virus** | **Reference** |
| --- | --- | --- |
| Phycocyanobilin | Influenza virus | Chen et al., 2016 |
| Quercetin | Hepatitis C | Bachmetov et al., 2012 |
| Riboflavin | Human foreskin fibroblast-vesicular stomatitis virus | Jamison et al., 1989 |
| Cyanidin | H1N1 influenza virus | Kannan & Kolandaivel, 2018 |
| Daidzein | Dengue virus type-2 | Zandi et al., 2011 |
| Genistein | Herpes b virus | LeCher et al., 2019 |
| Catechin | Influenza virus | Song, Lee & Seong, 2005 |
| Resveratrol | MERS-CoV | Lin et al., 2017 |
| Curcumin | HSV-1 | Zandi et al., 2010 |
| Astaxanthin | HSV-1 | Santoyo et al., 2012 |
| β-carotene | HSV-1 | Santoyo et al., 2012 |
| Capsaicin | HSV-1 & 2 | Hafiz et al., 2017 |
| Gingerol | Human respiratory syncytial virus | San Chang et al., 2013 |
| Vanillin | H1N1 influenza virus | Hariono et al., 2016 |
| Eugenol | Human herpes virus | Benencia & Courreges, 2000 |
| Thymol | HSV-1 | Lai et al., 2012 |

**Figure legends:**

**Fig. S1:** Initial screening of phytochemicals (selected based on docking score and our availability) by in vitro enzymatic assays

**Fig. S2:** 3D binding pocket of CoVs M^pro^ enzymes with top model Phycocyanobilin (cyan color) and surrounding active site amino acid residues (yellow color) with in 3 Å; remaining amino acid residues shown as cartoon, polar interactions are represented as magenta color.

**Fig. S3:** 3D binding pocket of CoVs PL^pro^ enzymes with top model Phycocyanobilin (cyan color) and surrounding active site amino acid residues (chain A - yellow color, chain B – orange color) with in 3 Å; remaining amino acid residues shown as cartoon, polar interactions are represented as magenta color.

**Fig. S4:** 3D binding pocket of SARS-CoV-2 M^pro^ with top model other Phycobilins (cyan color) and surrounding active site amino acid residues (yellow color) with in 3 Å; remaining amino acid residues shown as cartoon, polar interactions are represented as magenta color

**Fig. S5:** 3D binding pocket of SARS-CoV-2 PL^pro^ with top model other Phycobilins (cyan color) and surrounding active site amino acid residues (chain A - yellow color, chain B – orange color) with in 3 Å; remaining amino acid residues shown as cartoon, polar interactions are represented as magenta color.

**Fig. S1:**

**M^pro^ assay**

| Phytochemicals (µm) | % of Activity |
| --- | --- |
| P Control | 100 |
| I Control | 1.6797 |
| PC 50 | 62.9615 |
| PC 100 | 34.4875 |
| QC 75 | 105.031 |
| QC150 | 56.0673 |
| GS 150 | 93.5979 |
| GS 300 | 78.7214 |
| CD 150 | 97.5979 |
| CD 300 | 87.2142 |
| RF 150 | 98.9787 |
| RF 300 | 94.74 |

**PL^pro^ assay**

| Phytochemicals (µm) | % of activity |
| --- | --- |
| P Control | 100 |
| I Control | 3.13338 |
| PC 50 | 59.8587 |
| PC 100 | 29.2944 |
| GS 150 | 78.8146 |
| GS 300 | 65.5412 |
| RF 70 | 80.0179 |
| RF 140 | 66.6053 |
| QC 300 | 75.3308 |
| QC 600 | 73.5533 |

**Fig. S2:**

**SARS-CoV-1**

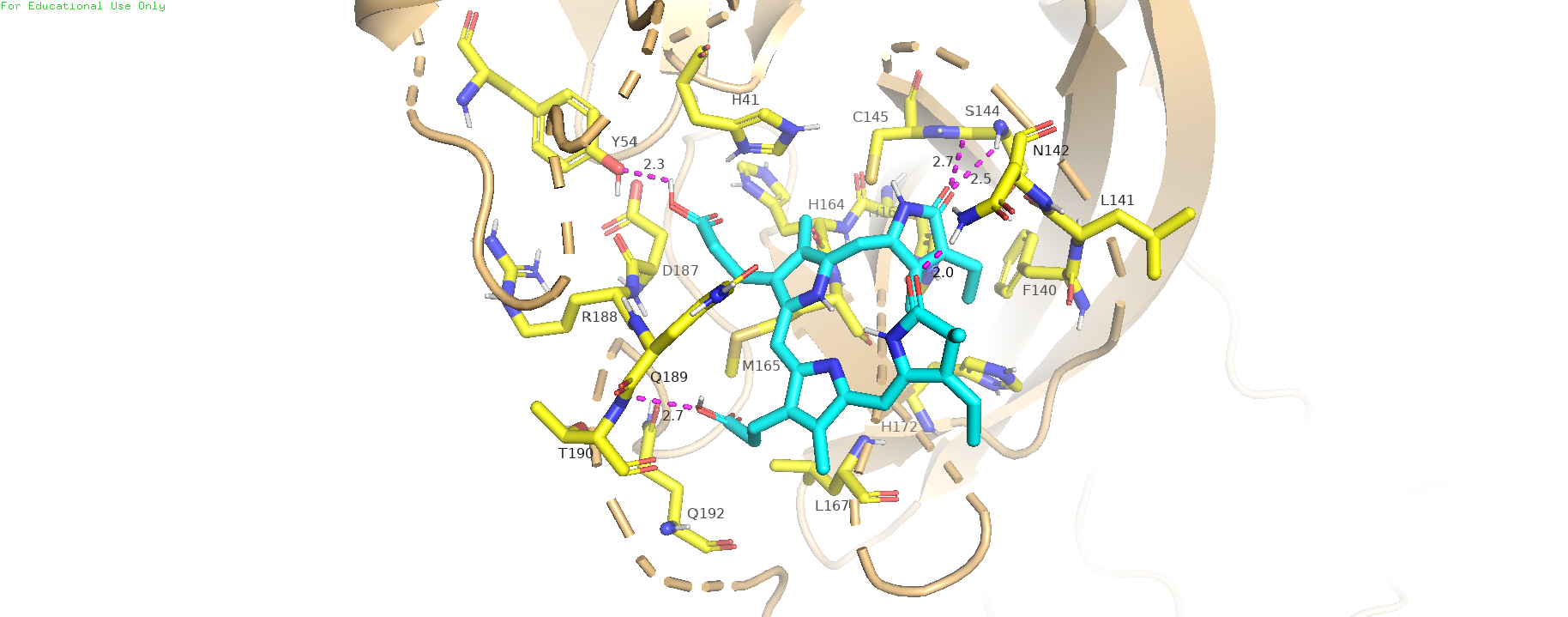

**MERS-CoV**

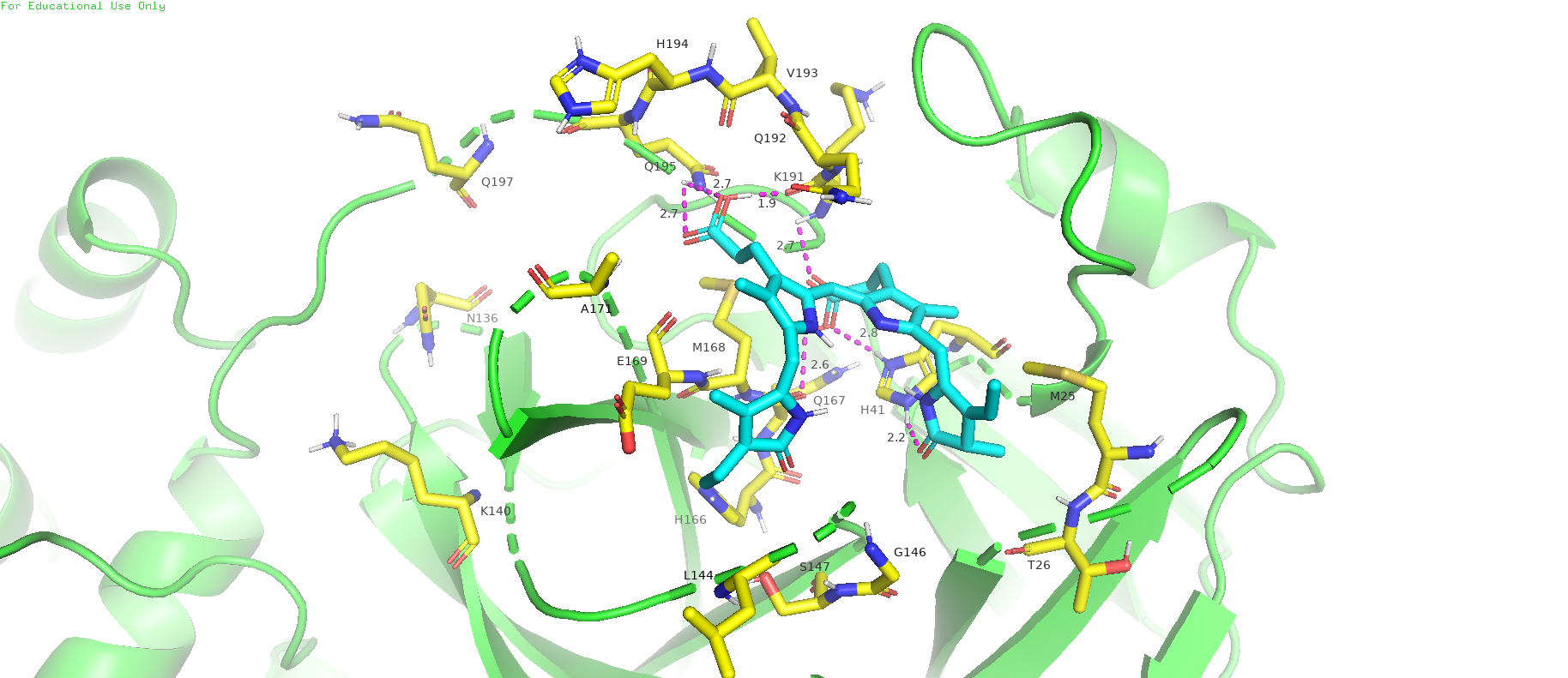

**HCoV-NL63**

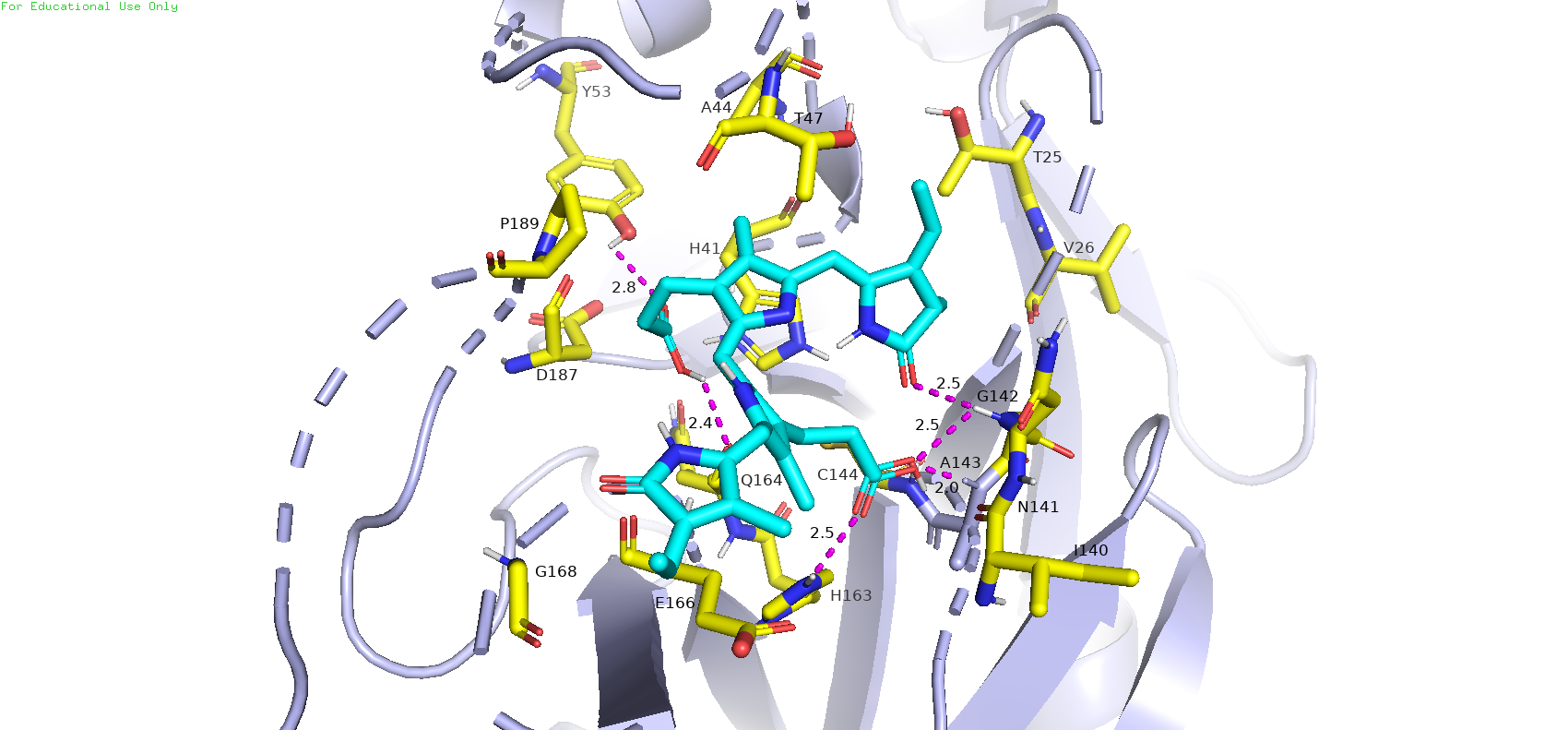

**HCoV-229E**

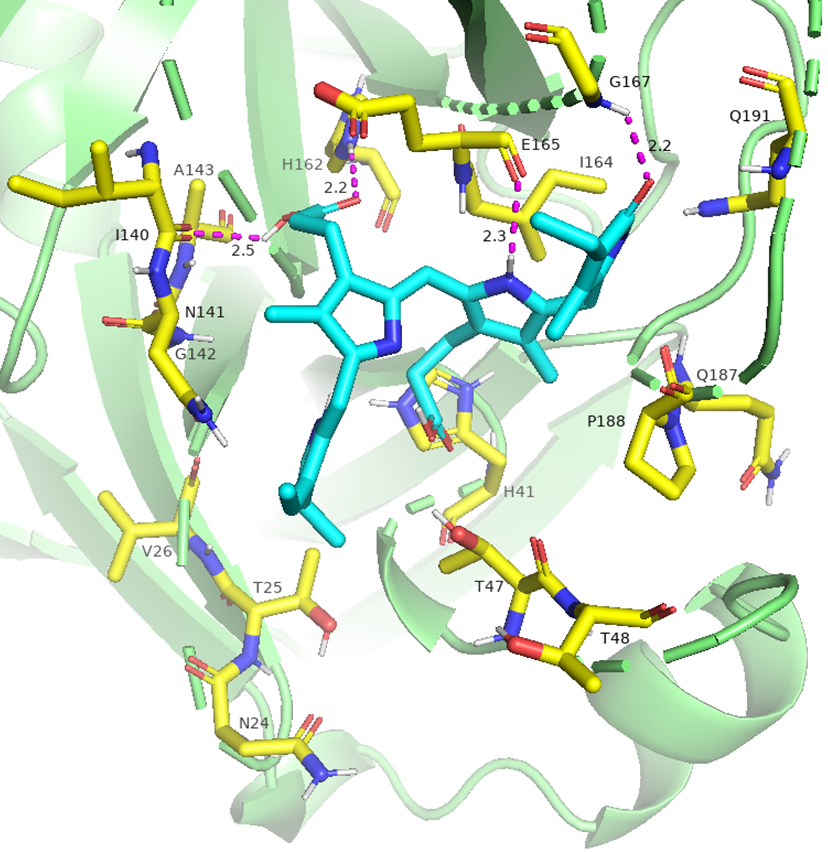

**HCoV HKU1**

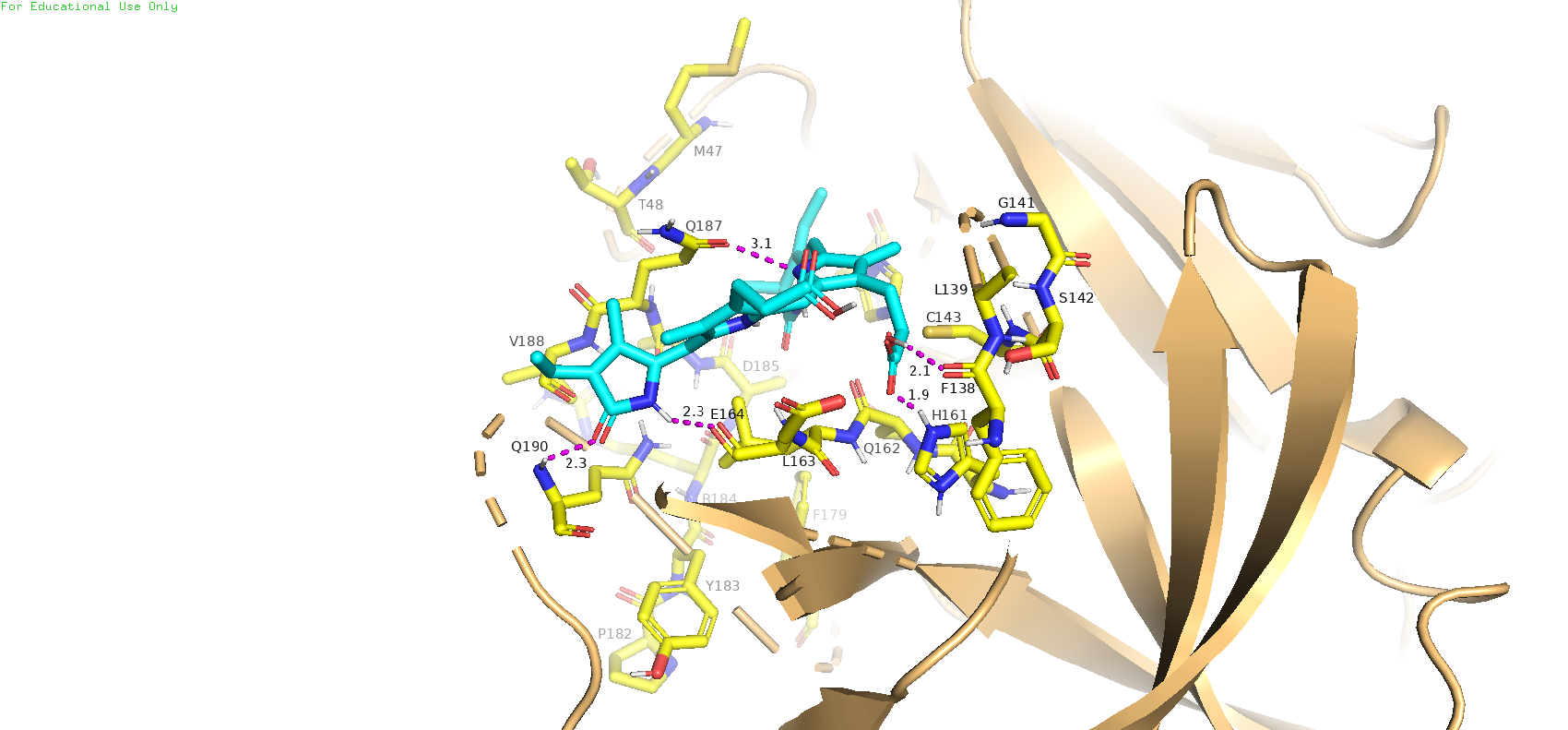

**MHV**

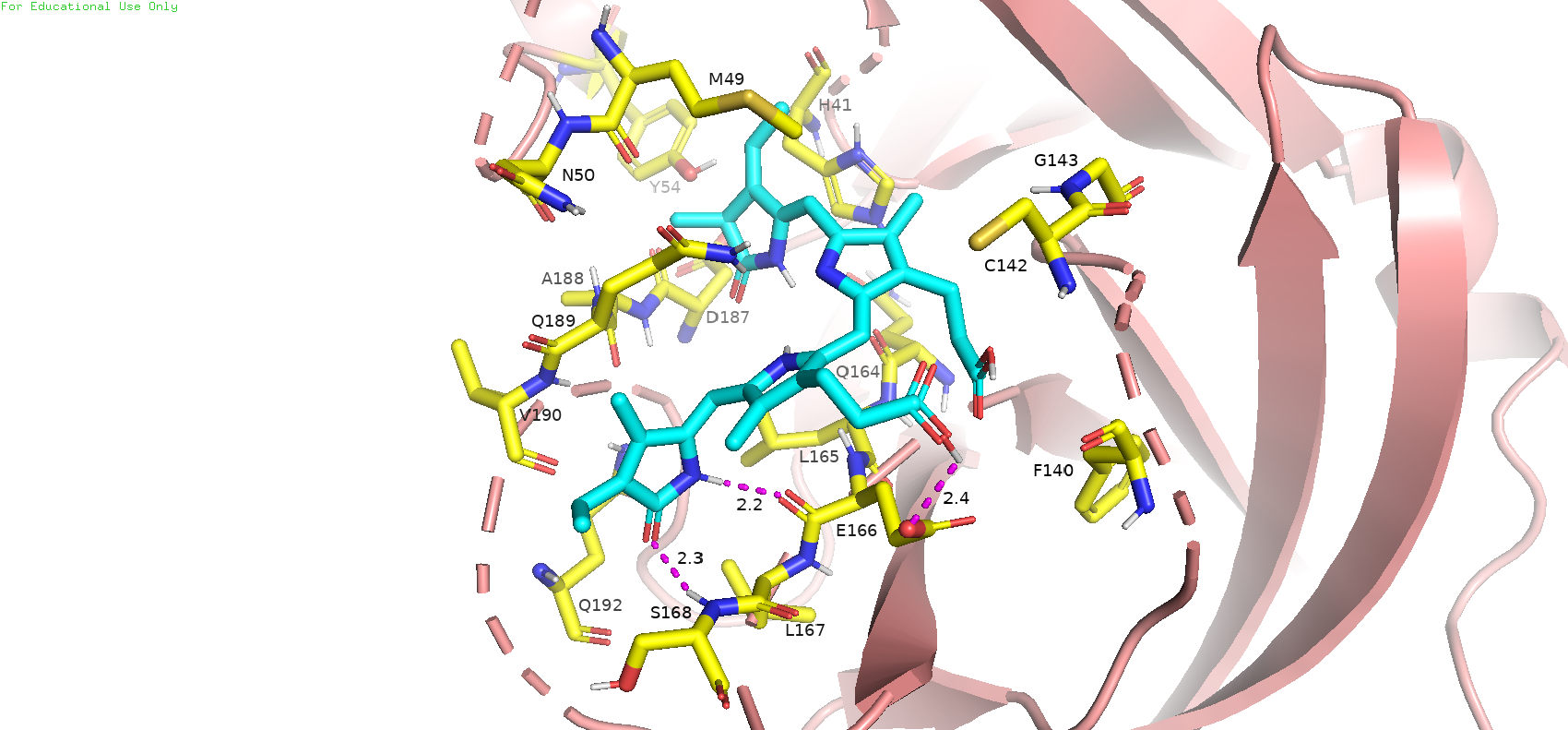

**TGEV**

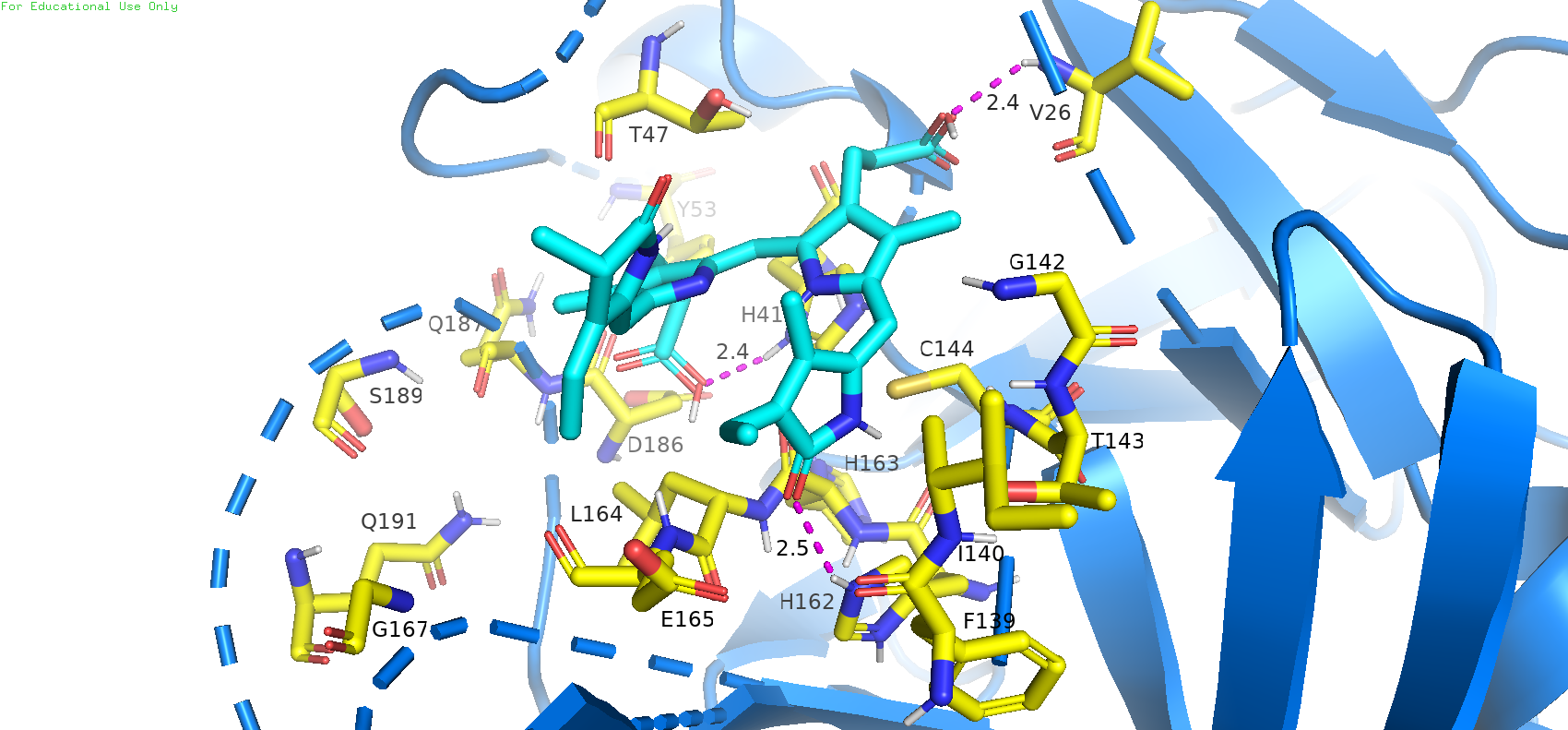

**FIPV**

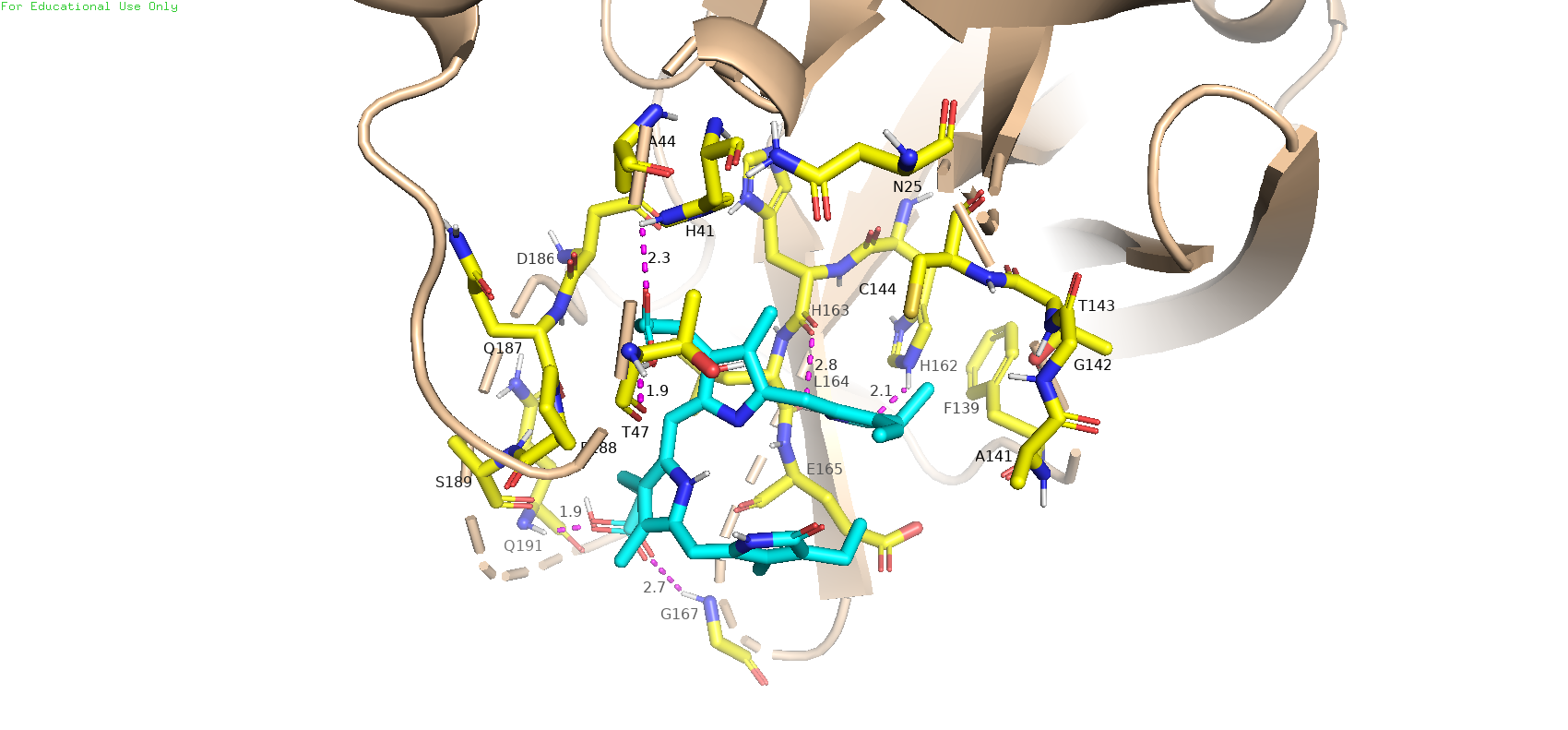

**IBV**

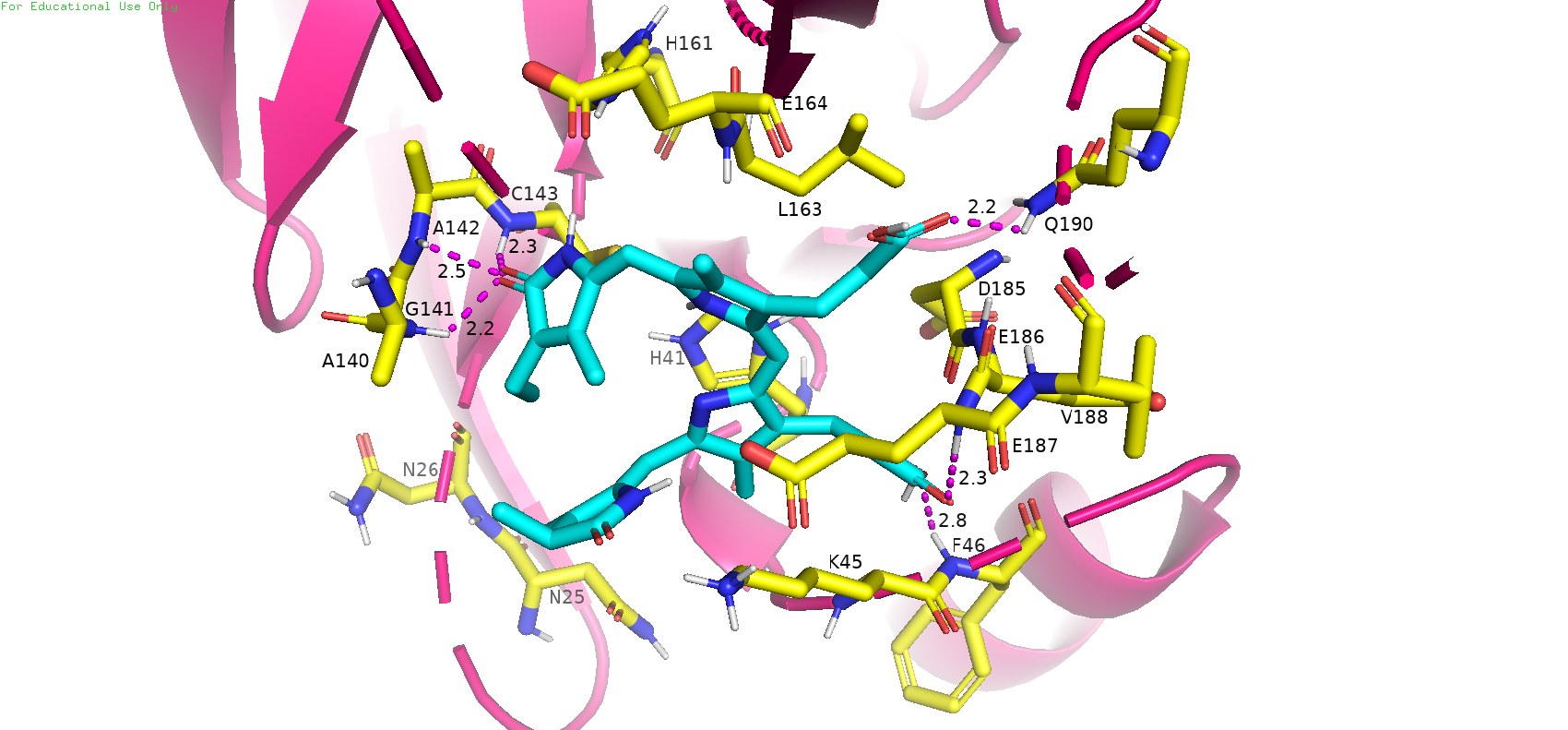

**Fig. S3:**

**SARS-CoV-1 (dimer)**

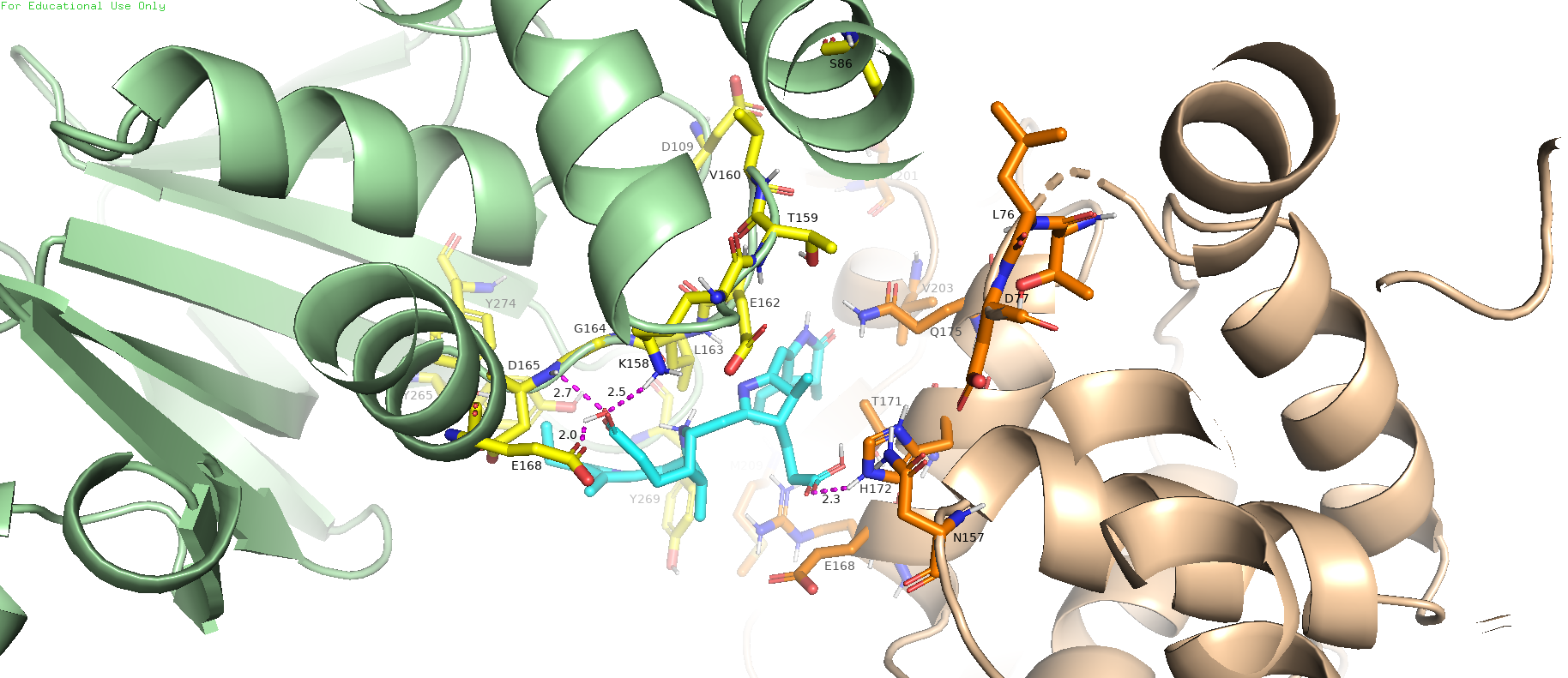

**SARS-CoV-1 (monomer)**

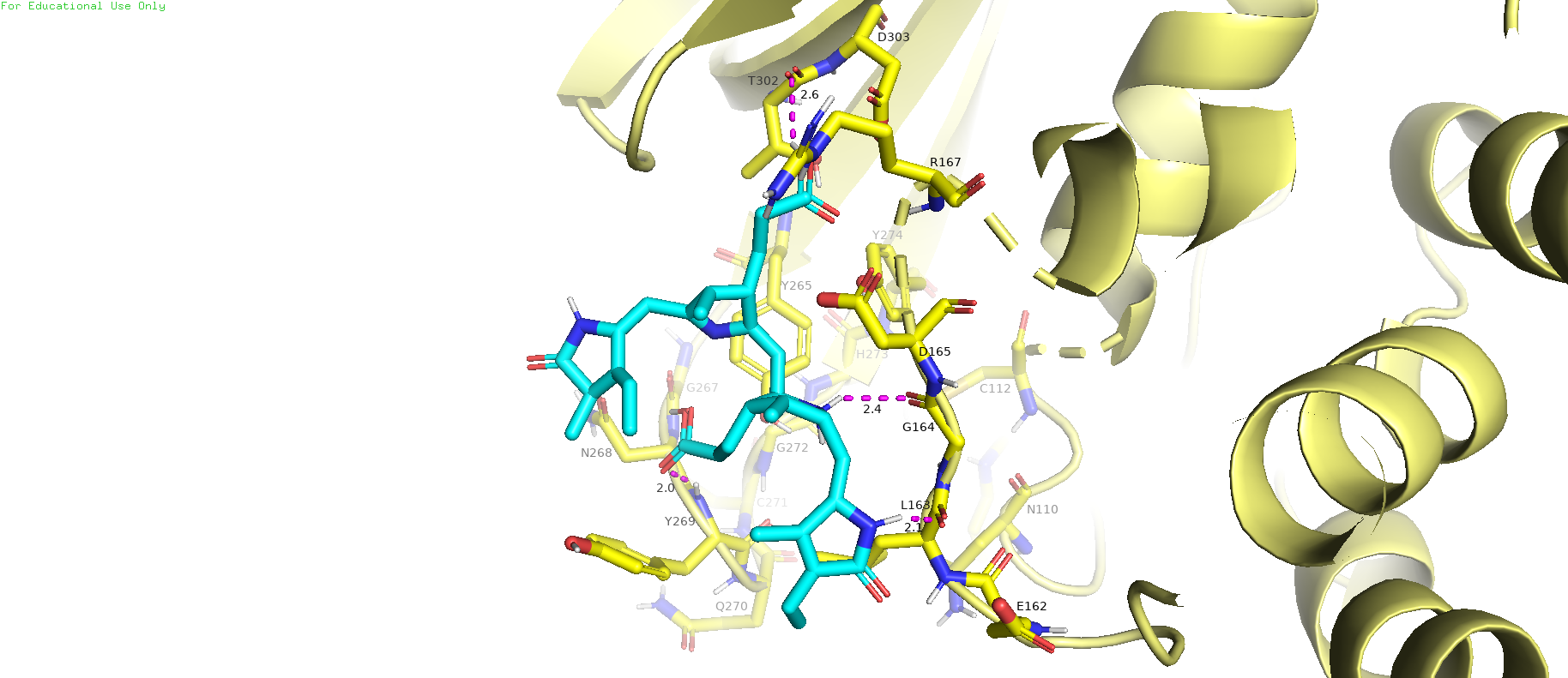

**SARS-CoV-2 (monomer)**

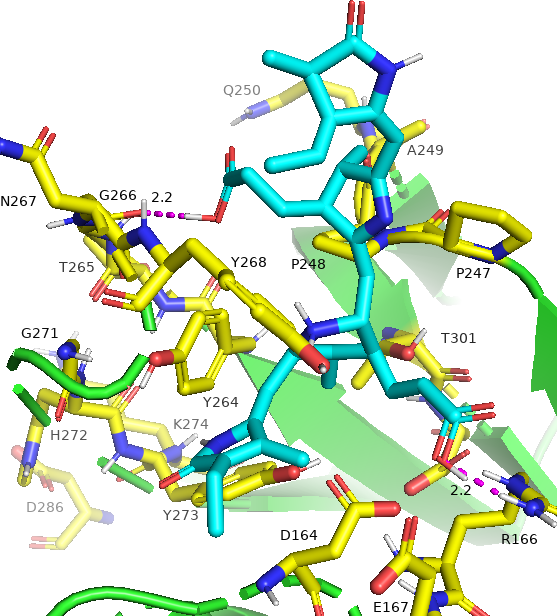

**MERS-CoV (monomer)**

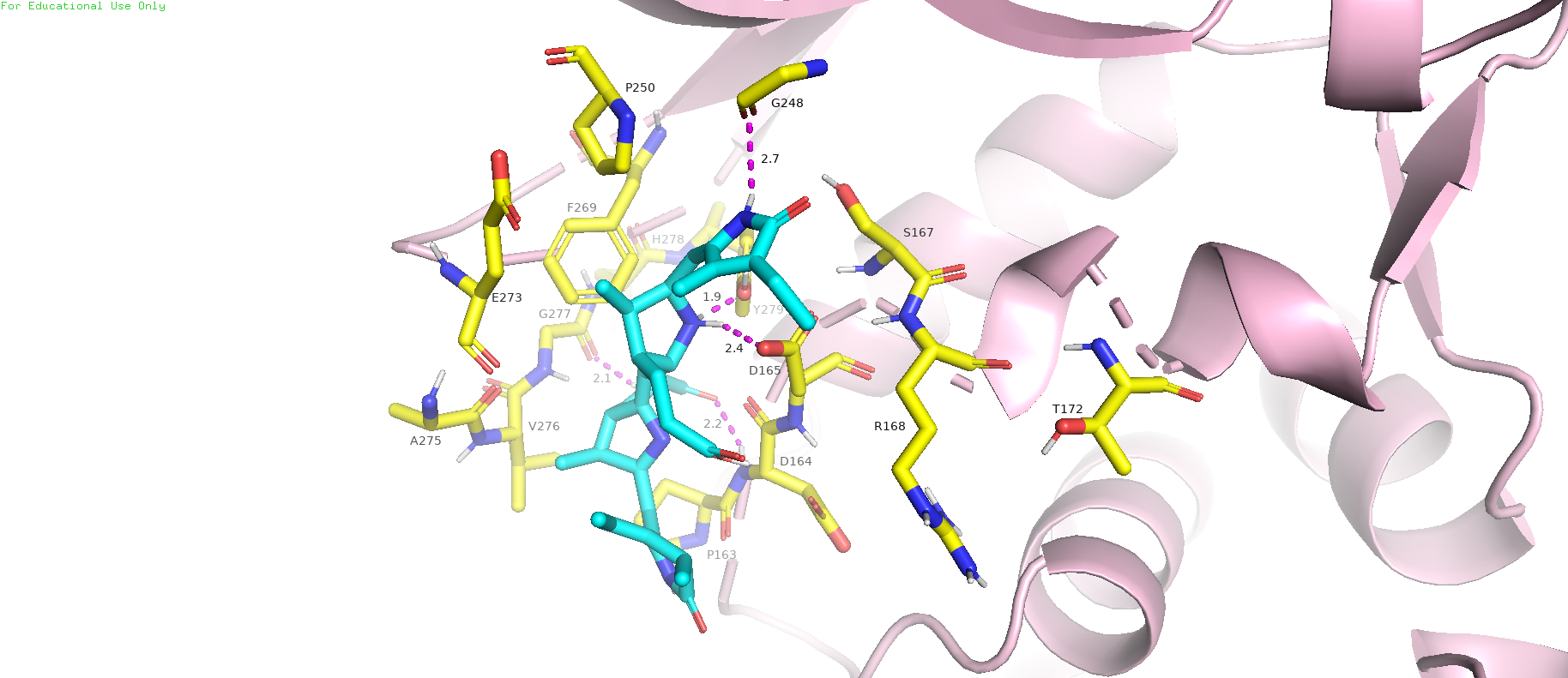

**TGEV (monomer)**

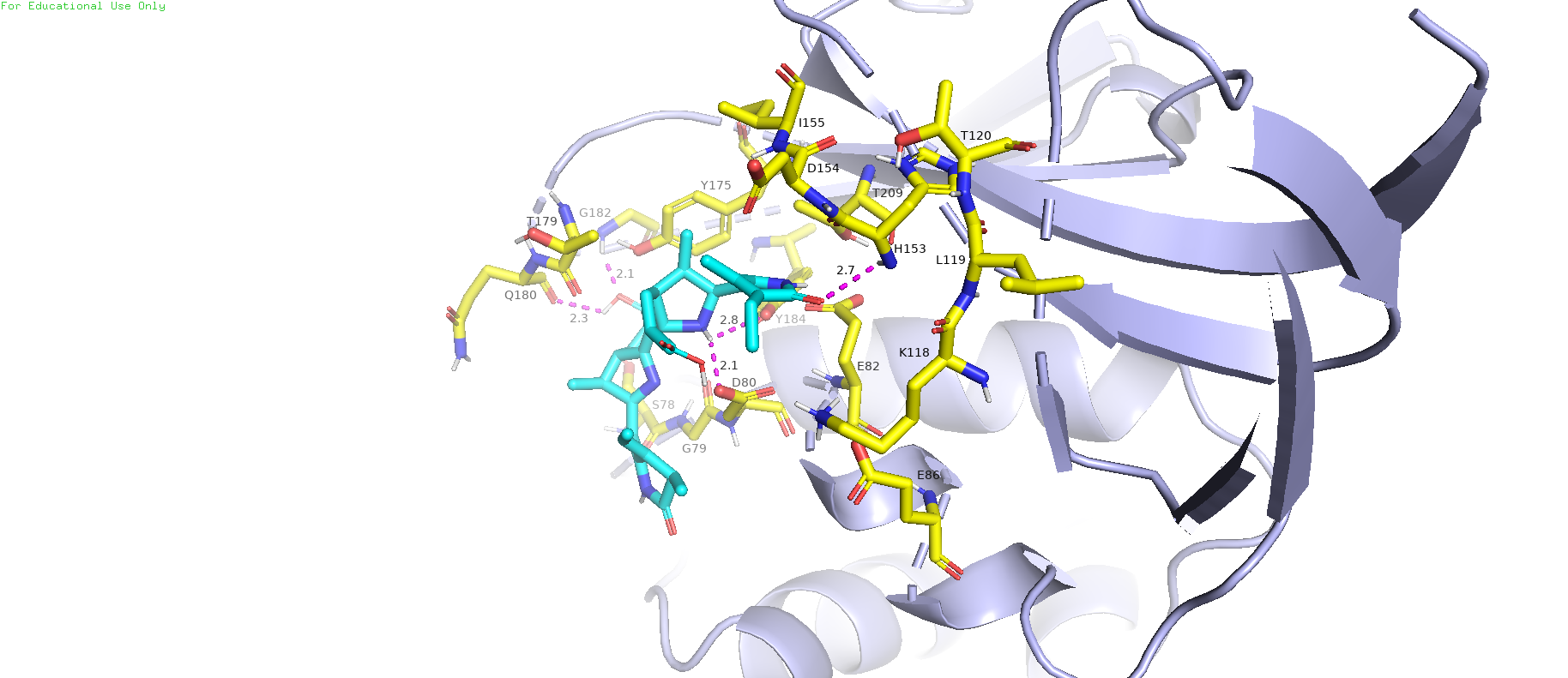

**IBV (monomer)**

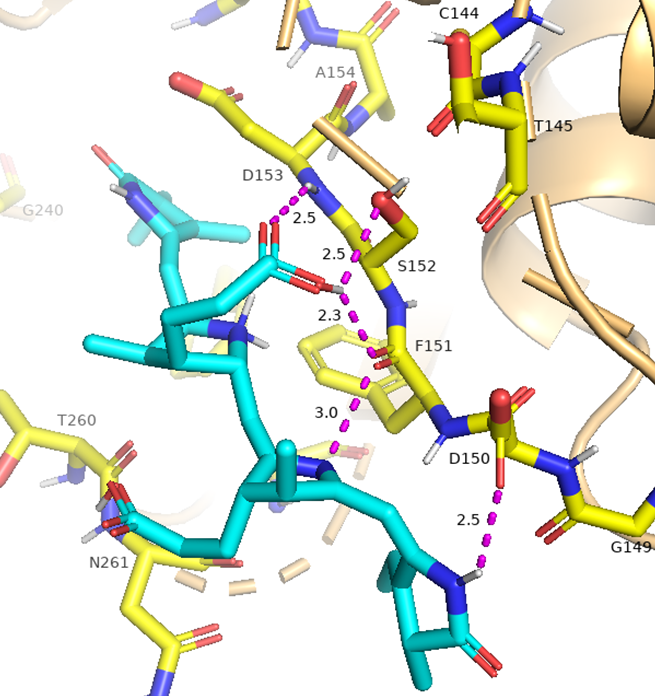

**Fig. S4:**

**Phycourobilin**

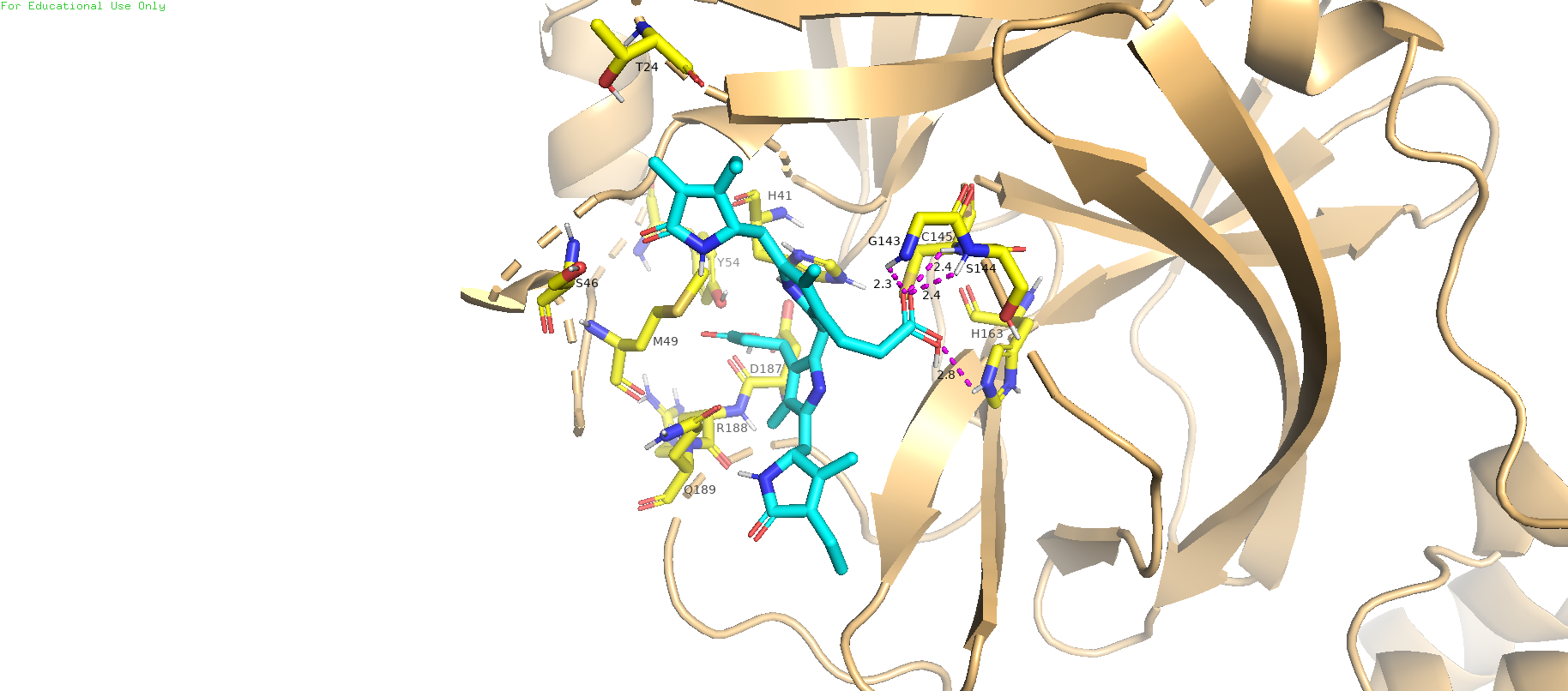

**Phycoerythrobilin**

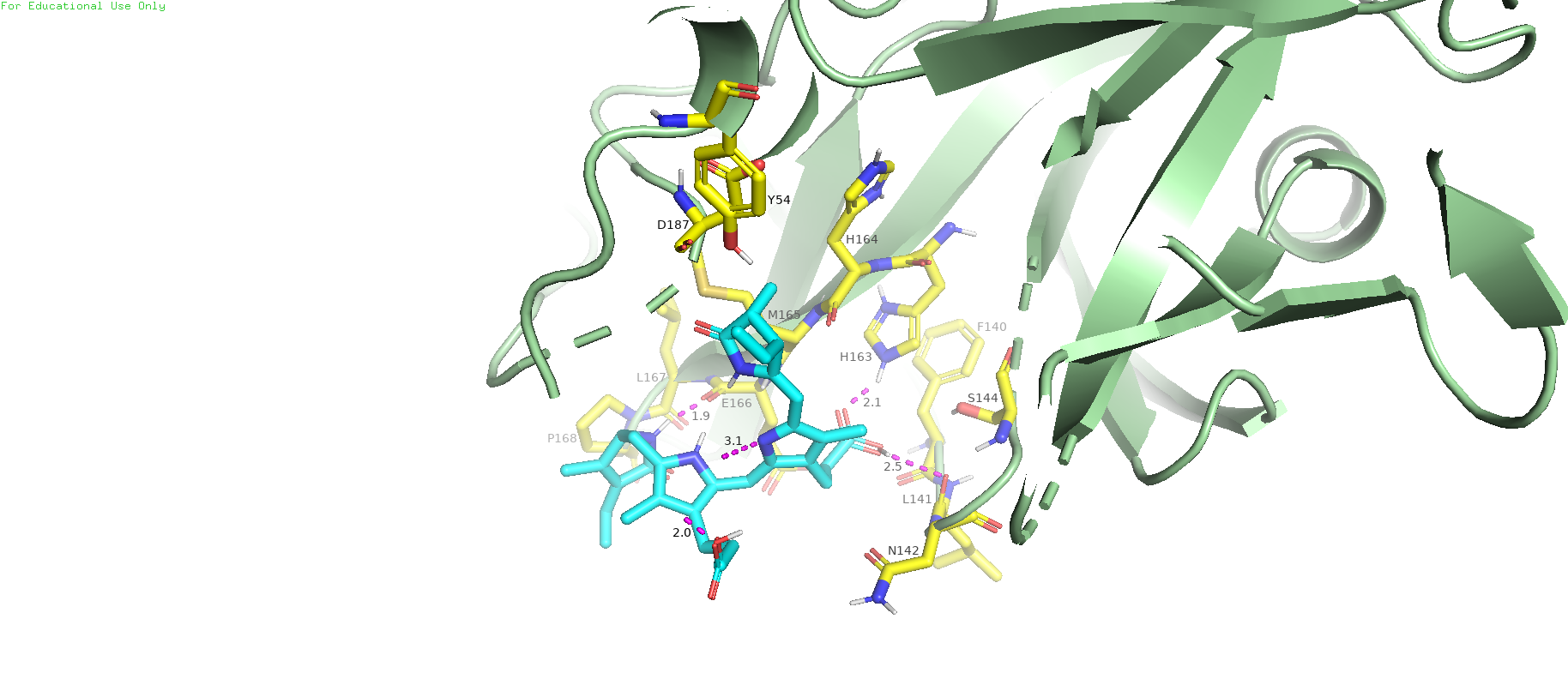

**Phycoviolobilin**

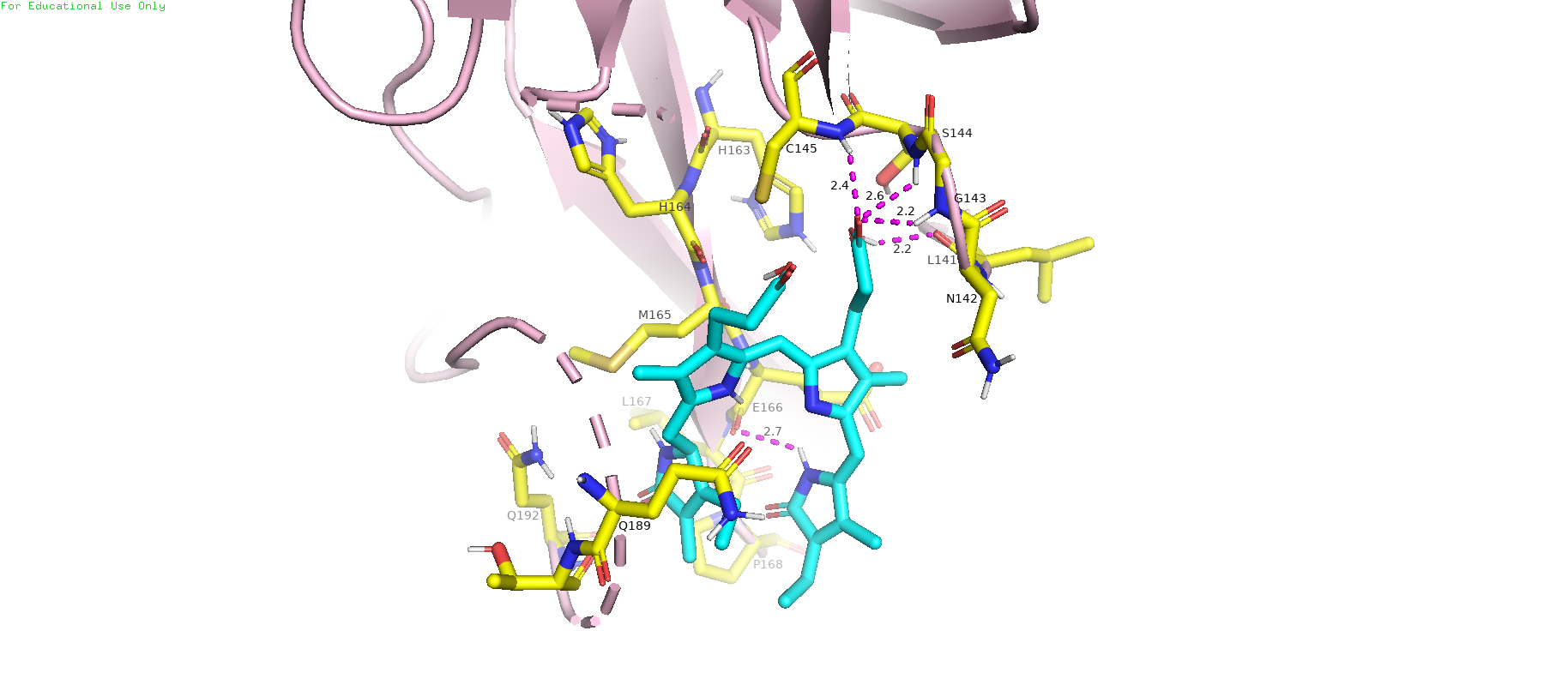

**Fig. S5:**

**Phycoerythrobilin**

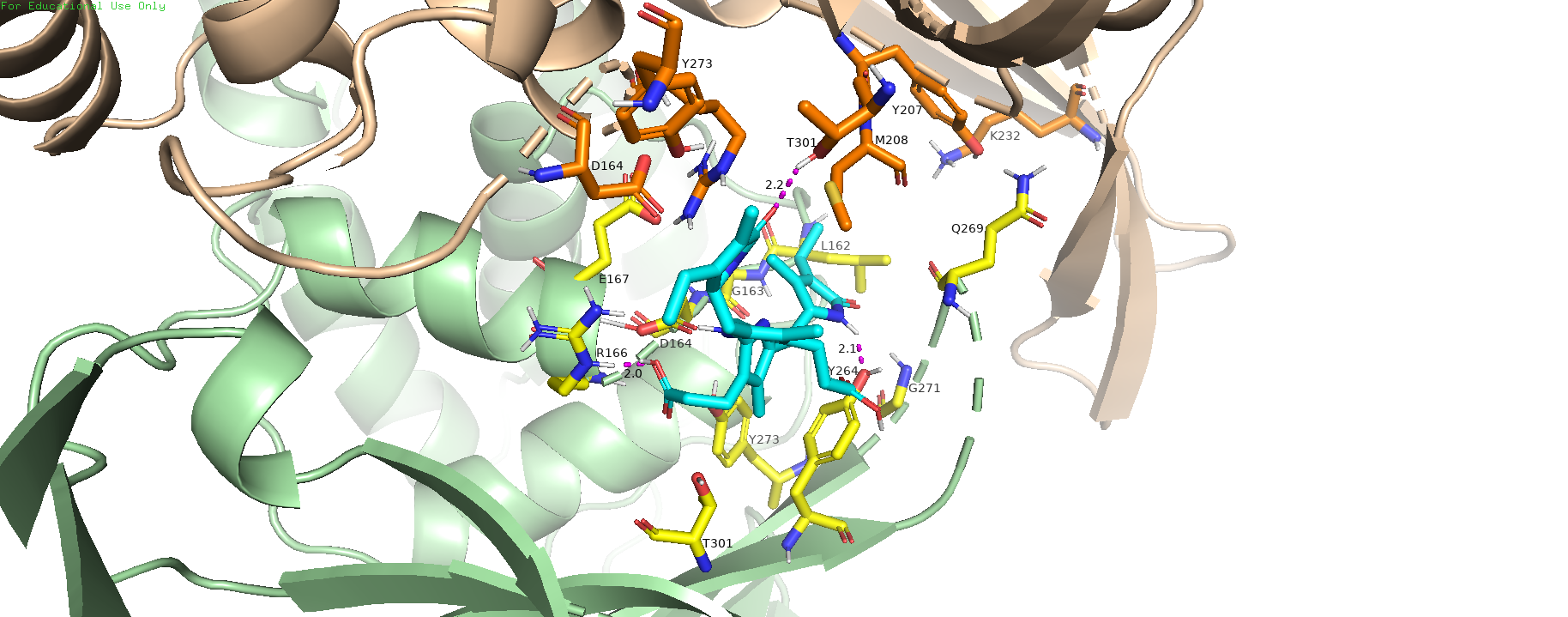

**Phycourobilin**

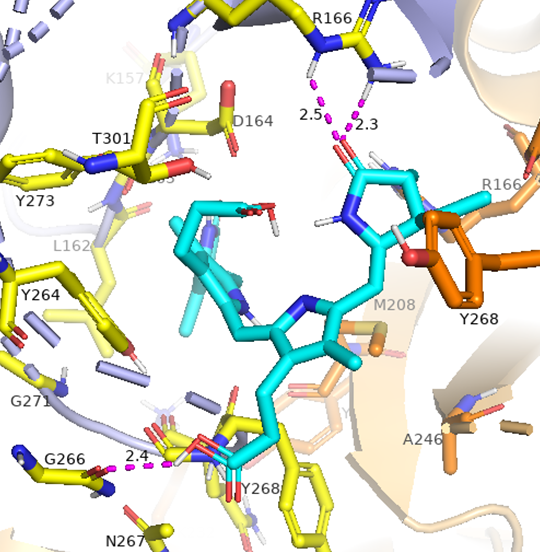

**Phycoviolobilin**

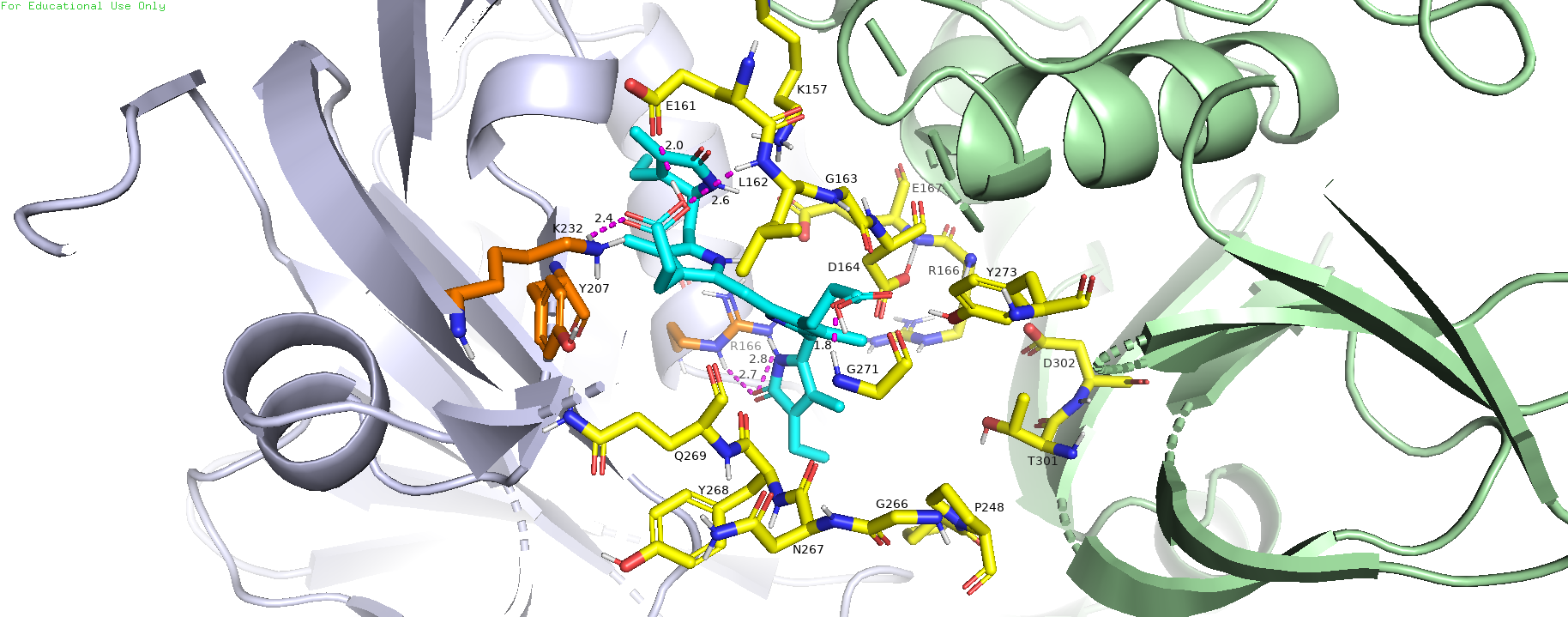

**References**

Bachmetov, L., Gal‐Tanamy, M., Shapira, A., Vorobeychik, M., Giterman‐Galam, T., Sathiyamoorthy, P., ... & Zemel, R. (2012). Suppression of hepatitis C virus by the flavonoid quercetin is mediated by inhibition of NS3 protease activity. *Journal of viral hepatitis*, *19*(2), e81-e88.

Benencia, F., & Courreges, M. C. (2000). In vitro and in vivo activity of eugenol on human herpesvirus. *Phytotherapy Research: An International Journal Devoted to Pharmacological and Toxicological Evaluation of Natural Product Derivatives*, *14*(7), 495-500.

Chen, Y. H., Chang, G. K., Kuo, S. M., Huang, S. Y., Hu, I. C., Lo, Y. L., & Shih, S. R. (2016). Well-tolerated Spirulina extract inhibits influenza virus replication and reduces virus-induced mortality. *Scientific reports*, *6*(1), 1-11.

Hafiz, T. A., Mubaraki, M., Dkhil, M., & Al-Quraishy, S. (2017). Antiviral activities of Capsicum annuum methanolic extract against herpes simplex virus 1 and 2. *Pak. J. Zool*, *49*, 251-255.

Hariono, M., Abdullah, N., Damodaran, K. V., Kamarulzaman, E. E., Mohamed, N., Hassan, S. S., ... & Wahab, H. A. (2016). Potential new H1N1 neuraminidase inhibitors from ferulic acid and vanillin: molecular modelling, synthesis and in vitro assay. *Scientific reports*, *6*(1), 1-10.

Jamison, J. M., Krabill, K., Hatwalkar, A., Jamison, E., & Tsai, C. C. (1990). Potentiation of the antiviral activity of poly r (AU) by xanthene dyes. *Cell biology international reports*, *14*(12), 1075-1084.

Kannan, S., & Kolandaivel, P. (2018). The inhibitory performance of flavonoid cyanidin-3-sambubiocide against H274Y mutation in H1N1 influenza virus. *Journal of Biomolecular Structure and Dynamics*, *36*(16), 4255-4269.

Lai, W. L., Chuang, H. S., Lee, M. H., Wei, C. L., Lin, C. F., & Tsai, Y. C. (2012). Inhibition of herpes simplex virus type 1 by thymol-related monoterpenoids. *Planta medica*, *78*(15), 1636-1638.

LeCher, J. C., Diep, N., Krug, P. W., & Hilliard, J. K. (2019). Genistein has antiviral activity against herpes b virus and acts synergistically with antiviral treatments to reduce effective dose. *Viruses*, *11*(6), 499.

San Chang, J., Wang, K. C., Yeh, C. F., Shieh, D. E., & Chiang, L. C. (2013). Fresh ginger (Zingiber officinale) has anti-viral activity against human respiratory syncytial virus in human respiratory tract cell lines. *Journal of ethnopharmacology*, *145*(1), 146-151.

Santoyo, S., Jaime, L., Plaza, M., Herrero, M., Rodriguez-Meizoso, I., Ibañez, E., & Reglero, G. (2012). Antiviral compounds obtained from microalgae commonly used as carotenoid sources. *Journal of applied phycology*, *24*(4), 731-741.

Song, J. M., Lee, K. H., & Seong, B. L. (2005). Antiviral effect of catechins in green tea on influenza virus. *Antiviral research*, *68*(2), 66-74.

Zandi, K., Ramedani, E., Mohammadi, K., Tajbakhsh, S., Deilami, I., Rastian, Z., ... & Farshadpour, F. (2010). Evaluation of antiviral activities of curcumin derivatives against HSV-1 in Vero cell line. *Natural product communications*, *5*(12), 1934578X1000501220.

Zandi, K., Teoh, B. T., Sam, S. S., Wong, P. F., Mustafa, M. R., & AbuBakar, S. (2011). Antiviral activity of four types of bioflavonoid against dengue virus type-2. *Virology journal*, *8*(1), 1-11.
